# CoupleVAE: coupled variational autoencoders for predicting perturbational single-cell RNA sequencing data

**DOI:** 10.1101/2024.03.05.583614

**Authors:** Yahao Wu, Jing Liu, Songyan Liu, Yanni Xiao, Shuqin Zhang, Limin Li

**Affiliations:** School of Mathematics and Statistics, Xi’an Jiaotong University, Xi’an, 710049, China; School of Mathematical Sciences, Fudan University, Shanghai, 200433, China

## Abstract

With the rapid advances in single-cell sequencing technology, it is now feasible to conduct in-depth genetic analysis in individual cells. Study on the dynamics of single cells in response to perturbations is of great significance for understanding the functions and behaviours of living organisms. However, the acquisition of post-perturbation cellular states via biological experiments is frequently cost-prohibitive. Predicting the single-cell perturbation responses poses a critical challenge in the field of computational biology. In this work, we propose a novel deep learning method called coupled variational autoencoders (CoupleVAE), devised to predict the post-perturbation single-cell RNA-Seq data. CoupleVAE is composed of two coupled VAEs connected by a coupler, initially extracting latent features for both controlled and perturbed cells via two encoders, subsequently engaging in mutual translation within the latent space through two nonlinear mappings via a coupler, and ultimately generating controlled and perturbed data by two separate decoders to process the encoded and translated features. CoupleVAE facilitates a more intricate state transformation of single cells within the latent space. Experiments in three real datasets on infection, stimulation and cross-species prediction show that CoupleVAE surpasses the existing comparative models in effectively predicting single-cell RNA-seq data for perturbed cells, achieving superior accuracy.

## 1 Introduction

Single-cell RNA sequencing (scRNA-Seq) technology has revolutionized biological and medical research by revealing the heterogeneity in a large number of individual cells. Breakthroughs in high-throughput scRNA-Seq data analyzing techniques have helped biologists understand the heterogeneity and transcriptomes change of specific cell types over a long period of life [1], explore the divergent developmental lineages of cell lines [2], quantify the pluripotency landscape of cell differentiation through cellular decomposition and dynamics [3, 4], and so on. Among the wide applications of scRNA-seq data, it is of great importance to study the dynamics of single cells in response to perturbations, such as virus infection [5], drug treatment [6, 7] or the knockout of genes [8, 9, 10], and the dynamic cellular differences for different species [11]. Such studies can help get a deep insight of the functions and behaviours of living organisms, and contribute to the discovery and development of combinatorial drugs. Identification of the marker genes associated with the perturbational dynamic differences can further enhance the understanding of the phenotypes from a cellular perspective [12]. However, due to the high cost of perturbational experiments [13, 14], computational modelling has been a necessary and effective way for predicting perturbation responses, which enables the generation of in silico scRNA-Seq data under specific conditions of interests.

While mechanistic models have been introduced in various studies for predicting single-cell perturbation responses [15, 16], such models usually have high computational costs and low precision in parameter estimation, and cannot explain the nonlinearity of the system. For example, a regularised linear model called MIMOSCA [8] is used to analyse the transcriptional effects of single-cell perturbations by modeling the expression level of each gene as a linear combination of the perturbation effects, which is not possible to model the complex nonlinear effects. In recent years, several deep generative methods for predicting single-cell perturbation responses have been proposed, especially using adversarial generative networks (GANs) and variational autoencoders (VAEs) [17, 18]. Particularly, the work in [19] designed a GAN-based generative model for simulating gene expression and predicting single-cell perturbations. Since GANs do not allow for mapping features into a latent space, another GAN-based method named scPreGAN [20] predicts single-cell by integrating autoencoder and GANs. However, it is often hard to train GANs with structured high-dimensional data, which limits the wide use of GANs for the single-cell perturbation prediction [21]. VAE-based generative models have been more popularly used [21, 22]. The method scGen [21] first learns the VAE latent feature differences between the perturbed and unperturbed cells and then maps the prediction in latent space to high-dimensional gene expression space. The method trVAE [22] uses CVAE [23] and maximum mean deviation to handle multiple perturbations. Both models consider the response of different cell types to the perturbation as the same and thus may ignore the different responses of different cell types to perturbations, which may lead to prediction bias. The MichiGAN model [24] combines VAEs and GANs to generate better samples while sampling from the untangled representation, where predicting drug response is one of its applications. However, this combined model usually has a complex structure, which may slow down the convergence rate and take a lot of time to train.

To overcome the limitations of the existing methods, in this work, we propose a deep generative model called coupling variational autoencoders (CoupleVAE) for predicting perturbation responses of scRNA-Seq data. CoupleVAE is a probabilistic graphical model that contains an inference process, a coupling process and a generative process. The main advantage of CoupleVAE is that the coupling process could learn the nonlinear mutual transformations between the controlled and the perturbed cells in latent space, which explains how the controlled cells respond to the perturbations. CoupleVAE not only enables predictions of SARS-COV-2 infection response of cells, IFN-*β* stimulation across different cell types, and it could also predict lipopolysaccharide perturbation of single cells from other species, with better performance than existing methods.

## 2 Results

### 2.1 Overview of the CoupleVAE

CoupleVAE aims to predict the responses of single cells measured by scRNA-Seq data after a specific type of perturbation. It takes the gene expression of controlled and perturbed cells as training data, and consists of an inference process (encoder), a coupling process (coupler) and a generative process (decoder). The inference process includes two independent encoders to obtain the lower-dimensional representation of cells under the control and the perturbation condition, respectively, in two separated latent spaces. Then the coupling process translates the two latent vectors using nonlinear transformations into the latent space of different conditions as a coupler. Finally, with the generative process by decoders, controlled cells and perturbed cells are reconstructed from the encoded vectors and the translated vectors in the same latent space, respectively.

The workflow of CoupleVAE is shown in Figure 1. Let **x**_*c*_ and **x**_*p*_ represent the gene expression for a controlled and perturbed cell, respectively. CoupleVAE first learns the posterior distributions *Q*_*ϕ*_(**z**_*c*_|**x**_*c*_) and *Q*_*ϕ*_(**z**_*p*_|**x**_*p*_) corresponding to the latent variables **z**_*c*_ and **z**_*p*_ for the controlled and perturbed cell by two encoders, respectively. The two latent variables are then mutually translated with conditional distributions 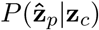 and 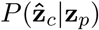 into the latent space of different conditions via coupling process. Finally, CoupleVAE generates gene expressions for controlled cells and perturbed cells by two decoders from translations 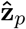 and 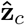, respectively.

**Figure 1:**
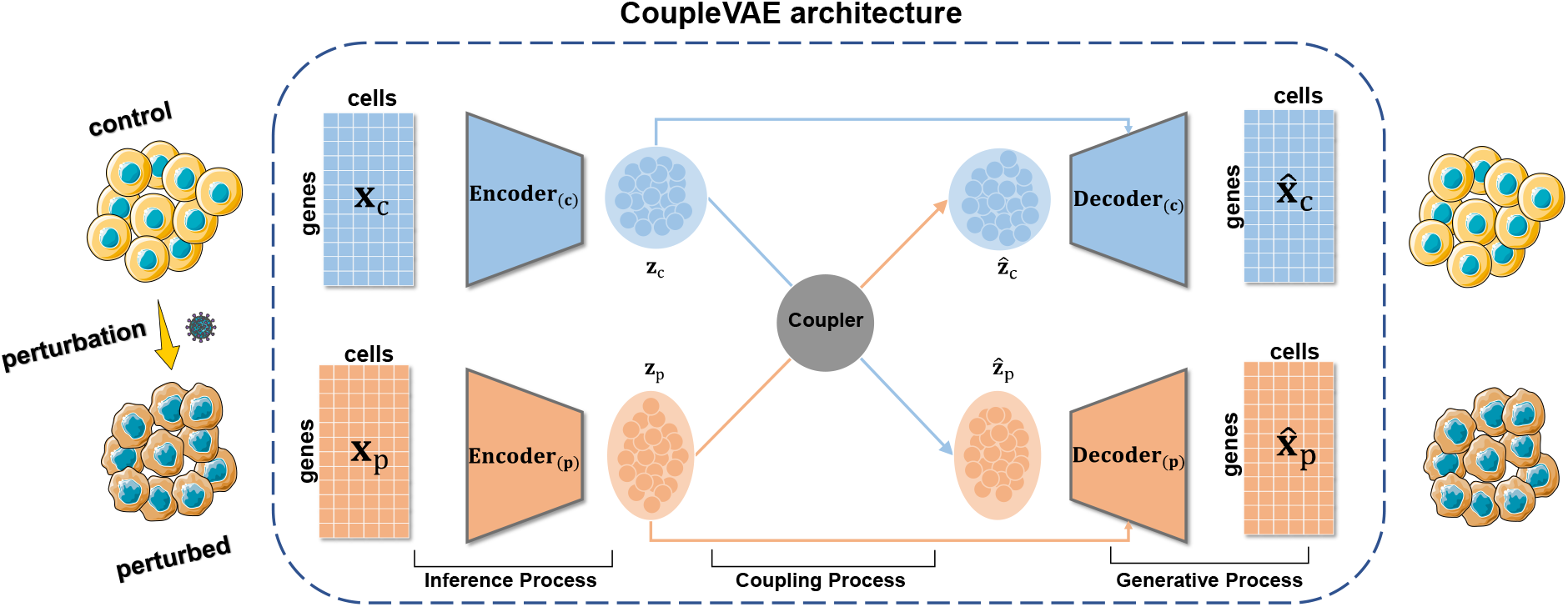
Workflow of CoupleVAE. CoupleVAE takes the gene expression of controlled cells (**x**_*c*_) and perturbed (**x**_*p*_) cells as input. The whole procedure includes three processes. Inference process: The latent representations **z**_*c*_ and **z**_*p*_ of controlled cells (**x**_*c*_) and perturbed cells (**x**_*p*_) are obtained by two encoders. Coupling process: **z**_*c*_ is transformed to 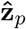 through a nonlinear mapping, and **z**_*p*_ is transformed to 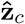 through inverse mapping by the coupler. Generative process: gene expression of the controlled cell 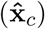 and the perturbed cell 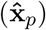 are reconstructed from **z**_*c*_, 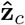 and **z**_*p*_, 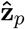 through the corresponding decoder. Components of this figure are created using Servier Medical Art templates, which are licensed under a Creative Commons Attribution 3.0 Unported License from https://smart.servier.com.

The experiments are done for three datasets including: the COVID-19 dataset with SARS-COV-2 infected cells, the PBMC dataset with IFN-*β* stimulation cells, and the LPS6 dataset with LPS6 stimulation for different species, with all dataset having their corresponding controlled cells. The following experiments show that CoupleVAE is able to predict the cells’ response to perturbation more accurately compared to the existing popular methods, and it enables cross-species perturbation prediction for an unseen species.

### 2.2 CoupleVAE accurately predicts response to SARS-COV-2 infection

This dataset contains cells from the COVID-19 dataset summarised by Lotfollahi *et al*. [25](Figure 2a), where the samples are collected from patients of different ages. We select six cell types in two conditions (control and severe COVID-19), and use the independently-and-identically distributed (i.i.d.) setting proposed by Bunne *et al*. [26](see Section 3) to predict the single cell responses to SARS-COV-2 infection. For example, in predicting macrophages’ response to perturbation, the training data contains only the cells of type macrophages. To evaluate the performance of CoupleVAE, the scRNA-Seq data for all six cell types after infection with SARS-CoV-2 are predicted and compared with the real perturbational data. We compare CoupleVAE with three single-cell perturbation prediction methods: scPreGAN [20], trVAE [22] and scGen [21], and one machine learning method CVAE [23], which has recently been adapted to preprocessing, batch-correcting and differential testing of single-cell data [17].

**Figure 2:**
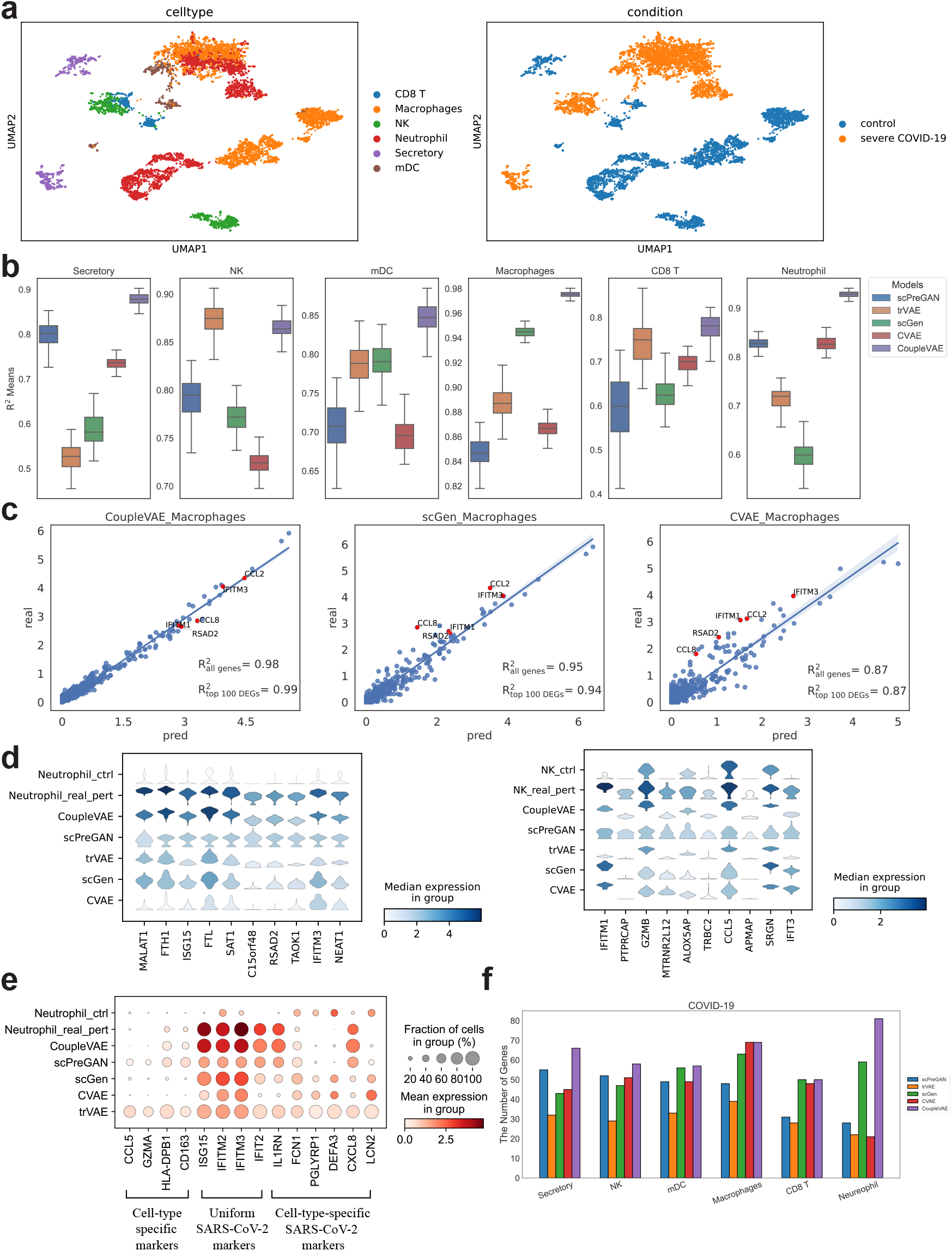
Single cell response prediction after SARS-CoV-2 infection. **a**. UMAP visualization of the distribution for cells of different types with and without SARS-COV-2 infection. **b**. Comparison of correlations between the real expression of all genes with SARS-COV-2 infection and the predicted using five methods. *R*^2^ denotes the square of the PCC between the means. **c**. Correlation comparison between the real mean values of 6000 genes’ expression in 248 macrophage cells with SARS-CoV-2 infection and those predicted by CoupleVAE, scGen and CVAE. **d**. Predicted gene distribution for 10 differentially expressed genes in neutrophil and NK cells after infected by SARS-CoV-2. The vertical axis shows the gene expression and distribution of neutrophil & NK cells in control, real perturbed and predicted by five different methods. **e**. Predicted mean gene expression comparison for the neutrophil cells with SARS-COV-2 infection. **f**. DEG identification results comparison. The number of genes represents the number of common genes between ground truth and predicted value among the top 100 DEGs.

We first calculate the correlations between the predicted and the real expression of all genes for each specific cell type, which are measured by the square of Pearson correlation coefficient (PCC) *R*^2^. Figure 2b and Supplementary Figure 1a report the boxplot of *R*^2^ for all genes and the top 100 deferentially expressed genes (DEGs), respectively. Results for the compared methods are also presented. Here, the DEGs are obtained by Scanpy [27] based on the controlled and real perturbational scRNA-Seq data. The results show that CoupleVAE could predict perturbational responses much more reliably than other methods except for NK cells, where though CoupleVAE achieves lower 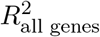 compared to scGen for the means of all genes’ expression, it obtains significantly higher correlation value 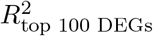 for top 100 DEGs (shown in Supplementary Figure 1a).

To examine the details of the predicted gene expression, we take a further look at the average of the predicted and real expression level for each gene in the cell type macrophages in Figure 2c and other cell types in the Supplementary Figure 2 and 3. Each point in the figure represents the average expression value of one gene across all cells in a given cell type. Compared to the two competitive methods scGen and CVAE, the expression level obtained from CoupleVAE tends to gather more closely to the real expression, and the 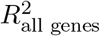 reaches 0.98, which is much higher than that of scGen (0.95) and CVAE (0.87). In particular, CoupleVAE predicts much more accurate expression values for the top 100 DEGs with 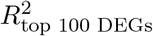 being 0.99, compared to that of scGen (0.94) and CVAE (0.87). The top five DEGs predicted by CoupleVAE also show a higher correlation with the real expression as marked in Figure 2c. We further examine whether the predicted gene expression distribution for each specific gene is close to the real distribution by looking at each of the top 10 DEGs in different cell types. Figure 2d shows the violin plots of the top 10 DEGs in neutrophil and NK cells, respectively, using the five different methods. CoupleVAE achieves more similar expression distributions for most of these genes in these two cell types, which implies that CoupleVAE could predict perturbational scRNA-Seq data with better accuracy. The violin plots for other cell types are also presented in Supplementary Figure 1b.

In addition, CoupleVAE can successfully predict the gene regulation patterns such as no regulation, shared regulation across all cell types, and specific regulation for one cell type. We use previous description of marker genes for three different gene regulation patterns: cell-type-specific markers corresponding to no regulation [28, 29, 30, 31, 32], uniform SARS-Cov-2 markers corresponding to shared regulation [31], and cell-type-specific SARS-Cov-2 markers corresponding to specific regulation [31]. Figure 2e and Supplementary Figure 4 show that the gene regulation pattern after infection could be captured accurately by CoupleVAE. Cell-type-specific markers show no expression changes before and after perturbation such as gene CCL5 and HLA-DPB1. Uniform SARS-Cov-2 markers change similarly for all cell types, for example, ISG15, IFIM2, IFIM3, and IFIT2 are upregulated for all cell types after perturbation. Cell-type-specific SARS-Cov-2 markers change differently for different cell types, for example, SARS-Cov-2 gene DEFA3 is downregulated only for neutrophils cells, and LCN2 is downregulated for neutrophils cells and upregulated in secretory cells. These results imply the ability of CoupleVAE in accurately predicting the gene regulation patterns after infection.

Finally, CoupleVAE is further evaluated by comparing the identified DEGs. The DEGs are identified using the methods in Scanpy [27] following the setting in scPreGAN [20]. That is, we take the top 100 DEGs between the controlled data and the real perturbational data as the ground truth, and identify the top 100 DEGs from the controlled data and the predicted data as the predicted DEGs. Then the number of the overlapping DEGs is computed. A higher percentage implies a more reliable model for predicting perturbational data. As shown in Figure 2f, among the five methods on the six cell types, CoupleVAE shows its better ability in identifying the ground truth DEGs. Particularly, CoupleVAE could identify more than 50 ground truth DEGs for all cell types, and even more than 80 DEGs for neutrophil cells among the top 100 DGEs.

### 2.3 CoupleVAE accurately predicts cell response for *IFN −β* treatment

We benchmark against a single-cell gene expression dataset containing 7217 *IFN −β* stimulated and 6359 controlled peripheral blood mononuclear cells (PBMCs) from eight different human lupus patients. Stimulation with *IFN−β* induced significant changes in the transcriptional profile of immune cells, which resulted in visible changes between control and stimulated cells. CoupleVAE and the four comparison models are applied to predict the single cell responses after IFN-*β* treatment. Similar to the strategy used for the COVID-19 dataset, we follow the i.i.d. settings for training and testing.

The performance of CoupleVAE is first evaluated by computing the square of PCCs *R*^2^ between the predicted and the real gene expression levels of either all genes or the top 100 DEGs, as shown in Figure 3a and Supplementary Figure 5a, respectively. CoupleVAE can obtain the highest or competitive *R*^2^ values for all cell types, which implies that it could simulate reliable gene expression levels after treatment perturbation.

**Figure 3:**
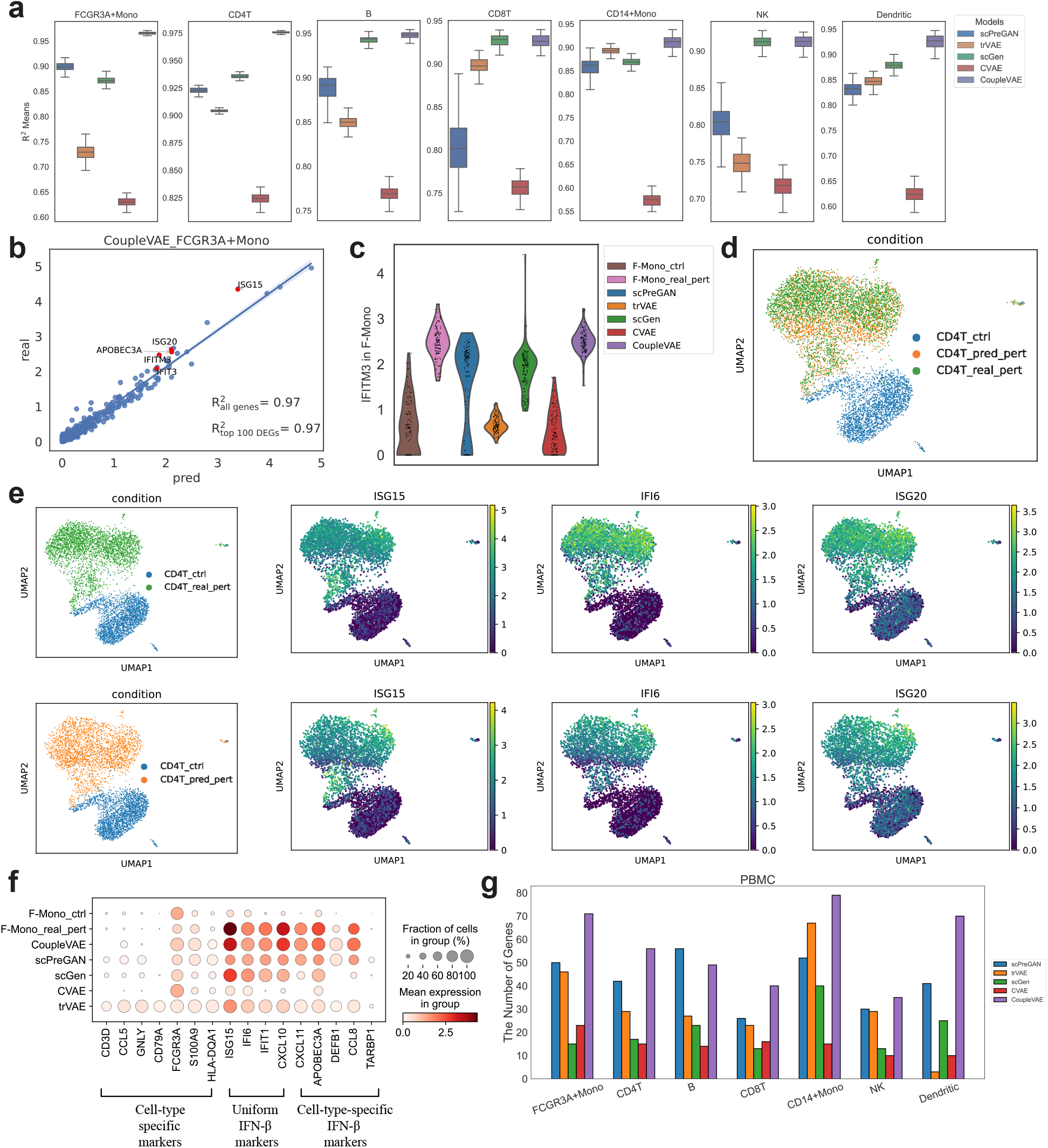
Single cell response prediction for *IFN − β* treatment. **a**. Boxplot of correlations between the predicted gene expressions and real ones. *R*^2^ denotes the square of the PCC. **b**. The mean value of 6998 gene expressions in 137 FCGR3A+Mono predicted using CoupleVAE compared to the real expressions. **c**. Distribution of the most strongly changed gene IFITM3 in FCGR3A+Mono with *IFN −β* perturbation in control, real and predicted stimulated FCGR3A+Mono cells of CoupleVAE compared to other prediction models. **d**. UMAP visualization of controlled data, real perturbed data and predicted data using CoupleVAE for CD4T cells. **e**. UMAP visualization of the expression of three genes, ISG15, IFI6 and ISG20 in controlled data, real perturbed data and predicted data using CoupleVAE for CD4T cells. The left figure shows the pattern of all genes under different conditions. The upper line shows real perturbational data and the lower line shows predicted. **f**. Comparison of gene expression for control, real and predicted perturbation on the FCGR3A+Mono(F-Mono) from PBMC for five methods. **g**. Comparison of DEG identification with five models in PBMC. The number of genes represents the number of common genes between ground truth and predicted value in the top 100 DEGs.

The details of the predicted expression level for each gene are further explored by looking at the distribution compared to the real expression in each specific cell type. Figure 3b plots the mean of the predicted expression by CoupleVAE and the real expression for each gene in cell type FCGR3A+Mono, which shows that the dots are concentrated around the straight line. Particularly, among all the genes, the predicted of the top 5 DEGs lie closely to the real ones. This implies that CoupleVAE could produce gene expression close to the real expression. Results for other cell types are shown in Supplementary Figure 6 and 7. Furthermore, Figure 3c compares distribution of the predicted and real gene expression for gene IFITM3 (the top response gene to *IFN−β*) in cell type FCGR3A+Mono(F-Mono). Supplementary Figure 5b shows the violin plots for the other five top DEGs. The violin plots show that CoupleVAE predicts the expression distribution most similar to the real one.

CoupleVAE could predict the gene regulations responding to the treatment similar to those in the real treatment data. The visualization of CD4T cells by UMAP [33] before and after treatment in Figure 3d shows that the predicted perturbational scRNA-Seq data by CoupleVAE is similar to the real one, and Figure 3e further gives some examples where the predicted expression values of three genes (ISG15, IFI6, ISG20) show the same tendency of up-regulation as the real ones in CD4 cells. Supplementary Figure 8 and Figure 9 show the same results for the other six cell types. Supplementary Figure 10 shows the results of the four comparison methods on seven cell types. Figure 3f and Supplementary Figure 11 show how the different methods predict regulation styles for cell-type-specific markers, uniform IFN-*β* markers and cell-type-specific IFN-*β* markers for F-mono cells [34]. CoupleVAE could predict the regulation pattern better than the other methods. For example, in F-Mono cells, scGen fails to identify the upregulation for some uniform cell-specific IFN-*β* markers DEFB1 and CCL8 after treatment, CVAE misses the upregulation of cell-specific IFN-*β* markers such as IFIT1, CXCL20, and CXCL11, trVAE predicts all genes as upregulation. Figure 3g finally gives the number of common DEGs between the top 100 predicted DEGs obtained by predicted perturbational scRNA-seq and top 100 ground truth DEGs obtained by real scRNA-seq data. For six of the seven cell types, CoupleVAE could identify the highest number of true DEGs, which implies that the predicted perturbational scRNA-Seq captures the regulation information the best.

### 2.4 CoupleVAE accurately predicts cross-species perturbation responses

CoupleVAE could also be applied to predict cross-species perturbation responses, either under the same or different conditions. We collect the scRNA-Seq data [35] that studies the evolution of innate immunity programs of mononuclear phagocytes within different species, including pigs, rabbits, mice and rats. The primary bone marrow-derived cells are stimulated using lipopolysaccharide (LPS) for six hours. The data includes two groups: control phagocytes and phagocytes perturbed by LPS for 6h.

We refer to the task of predicting perturbations across species as out-of-distribution (o.o.d.), where unlike the previous two datasets, we expect different species respond to the stimulation differently. The cross-species perturbation prediction can be formulated as two scenarios. One is to train the prediction model using data from all species to predict the response of one of these species (Figure 4a), and the other is to train the model using one specific species to predict the perturbation response of another species (Figure 4e). Corresponding to these two scenarios, we set up two different training strategies: condition-CoupleVAE and species-CoupleVAE, where condition-CoupleVAE explains the differences of stimulation responses in all the considered species and species-CoupleVAE explains the transformations across different species.

**Figure 4:**
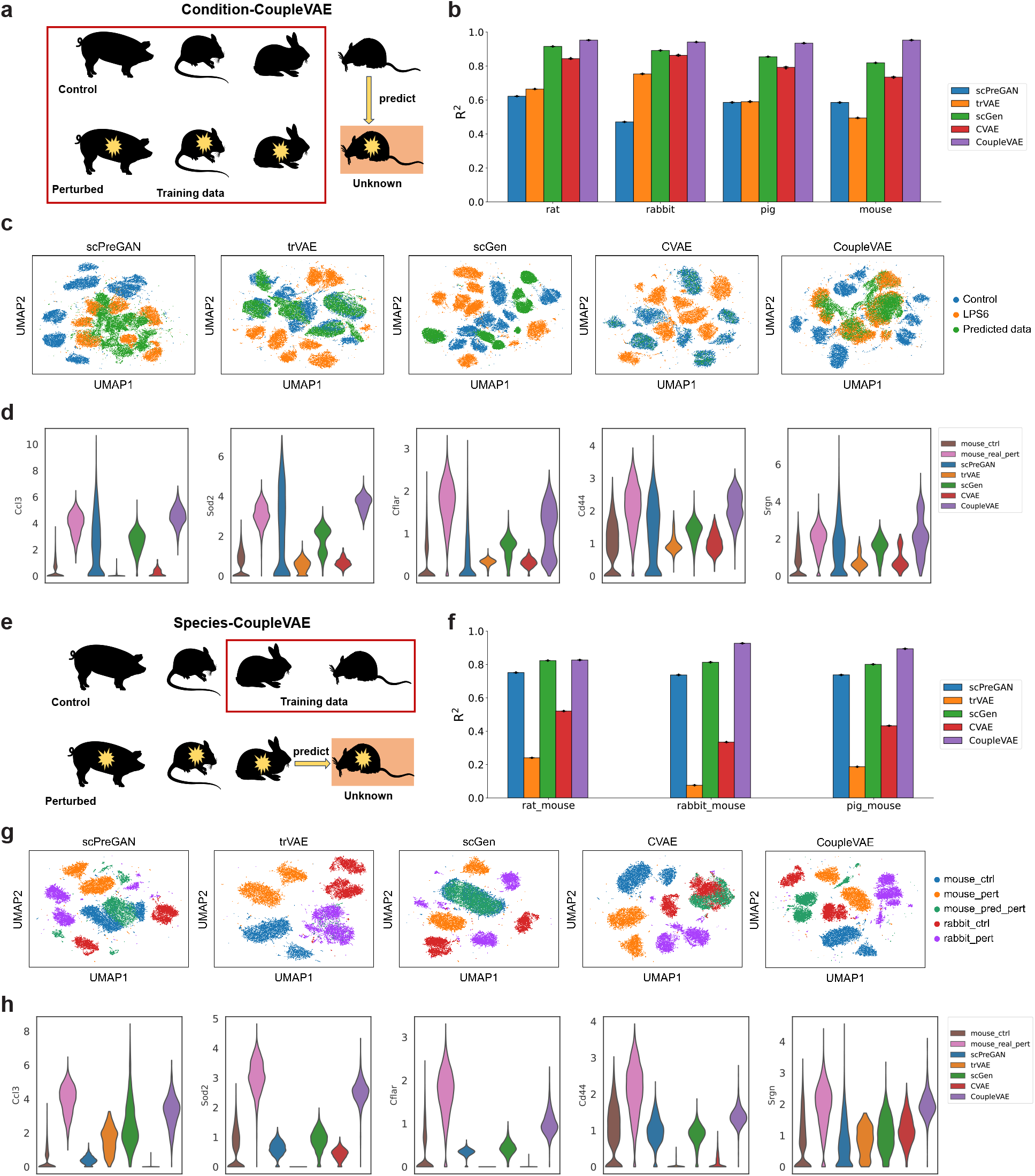
Cross-species single cell response prediction. **a**. The LPS dataset contains four species: rat, rabbit, pig and mouse. The first row represents the controlled species, and the second row is the species perturbed with lipopolysaccharide (LPS) for six hours. Experimental setup of condition-CoupleVAE: data in red box are used for training to predict unseen LPS perturbed mouse cells from mouse controlled cells. **b**. Prediction comparison for rat, rabbit, pig and mouse cells perturbed by LPS from the corresponding cells in a healthy state, respectively. *R*^2^ for the average expression of all genes are compared between real and predicted cells from different models. **c**. UMAP visualization of controlled data, real perturbational data and predicted perturbed data using five methods for all species. **d**. Distribution of LPS-responsive genes (*Ccl3, Sod2, Cflar, Cd44, Srgn* [35]). **e**. Experimental setup of species-CoupleVAE: data in red box are used for training to predict unseen LPS perturbed mouse cells from LPS perturbed rat cells. **f**. Prediction of mouse cells’ responses to LPS from rat, rabbit and pig cells perturbed by LPS, respectively. **g**. UMAP visualization of controlled data, real perturbational data for mouse and rabbit, and predicted mouse perturbational data. **h**. Distribution of LPS-responsive genes(*Ccl3, Sod2, Cflar, Cd44, Srgn* [35]). Vertical axis: expression distribution for genes. Horizontal axis: control, real and predicted distribution by different models. Figure a and e were created using Servier Medical Art templates, which are licensed under a Creative Commons Attribution 3.0 Unported License; https://smart.servier.com.

Condition-CoupleVAE trains the model using both controlled and perturbed data for single cells from different species as shown in Figure 4a, and aims to predict the responses to perturbation for one specific species based on its own controlled data. For example, mouse controlled cells can be used as test data to predict the responses to LPS of mouse with the trained model. Figure 4b shows the results for different models in predicting the response of the four species to the perturbation. CoupleVAE achieves the highest *R*^2^ in all the four species. To further explore whether the predicted data from each method match the distribution of the real data, we combine the predicted data from the four species with the real data, and visualise the distribution using UMAP. As shown in Figure 4c, the predicted data do not overlap with the real perturbational data well using the four comparison methods, and the predicted data of trVAE and CVAE even overlap with the controlled data. In contrast, the predicted data of CoupleVAE is closer to the real data. This is due to the fact that CoupleVAE interconverts the distributions of the data under the two conditions in the latent space, which makes the generated data closer to the real data. More detailed explanations of the data distribution in the latent space are provided in the Supplementary Figure 13 and Supplementary Note 5. We then select six DEGs that are differentially expressed before and after the mouse is perturbed by LPS, and study their predicted distribution using all five methods. For gene *Ccl3, Sod2, Cflar*, CoupleVAE predicts the entire distribution well. For some other genes (*Cd44, Srgn*), their distributions are closer to the true distribution compared to other methods. The predicted gene distributions associated with LPS perturbations for the other three species are shown in the Supplementary Figure 14.

Species-CoupleVAE is to predict the perturbation responses of one species based on the perturbational data from another species and the controlled data of both species. As shown in Figure 4e, the training data are rabbit and mouse cells in the control state, and the model predicts mouse cells perturbed by LPS from rabbit cells perturbed by LPS. Figure 4f demonstrates the *R*^2^ for the perturbational mouse cells predicted by different models from the other three species. CoupleVAE gives much higher *R*^2^ than all the other four methods. Figure 4g shows UMAP plots of the predicted perturbational data of mouse cells from rabbit cells. Each plot contains rabbit and mouse cells in the control and LPS states, as well as the predicted expression for mouse cells perturbed by LPS for the five methods. Among them, the predictions of scGen and trVAE overlap with the mouse cells in the control state. For CVAE, its predicted data partially overlaps with the rabbit cells in the control state. As for scPreGAN, the predicted data still partially overlaps with the mouse cells in the control state. The predicted data of CoupleVAE has no overlap with rabbit and mouse cells in control state, and they are closer to the real data distribution. Figure 4h is similar to Figure 4d, where the same genes are picked to study the prediction of gene distribution by species-CoupleVAE(training data is Figure 4e). CoupleVAE predicts the distribution of the above mentioned genes better.

## 3. Methods

We formulate the perturbational single-cell RNA sequencing data prediction problem as follows. Let **x**_*c*_ and **x**_*p*_ be the vectors representing the RNA sequencing data for a single cell before and after some perturbation. We aim to learn a predictive model *P* (**x**_*p*_|**x**_*c*_) based on the training pairs 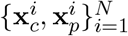.

### 3.1 Variational autoencoder

Variational autoencoder (VAE) [36] is a network structure to learn a generative process for multidimensional variable **x** through a latent variable **z** based on *N i*.*i*.*d*. samples 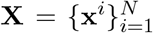 of **x**. It assumes a generative process which consists of generating **z**^*i*^ with a prior distribution *P*_*θ*_(**z**) and generating **x**^*i*^ from a conditional distribution *P*_*θ*_(**x**|**z**), where the parameters *θ* for the latent variables **z**^*i*^ are unknown. By introducing an encoder *Q*_*ϕ*_(**z**|**x**) to approximate the true intractable posterior *P*_*θ*_(**z**|**x**), the evidence lower bound (ELBO) of the marginal likelihood log *P*_*θ*_(**x**^*i*^) [37] can be written as:

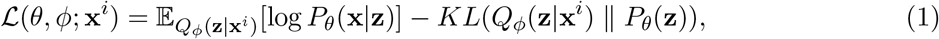

where 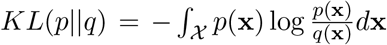 is the KL Divergence for two distributions *p* and *q*. The VAE jointly learns parameters *ϕ* in the encoder *Q*_*ϕ*_(**z**|**x**) and parameters *θ* in the decoder *P*_*θ*_(**x**|**z**) by maximizing the ELBO *ℒ* (*θ, ϕ*; **x**^*i*^), where the first term represents the reconstruction likelihood and the second term ensures that the learned distribution *Q*_*ϕ*_ is similar to the true prior distribution *P*_*θ*_.

In order to estimate the ELBO based on the given data points, the VAE takes prior of **z** as *P*_*θ*_(**z**) = *𝒩* (**z**; **0, I**), and assumes the variational distribution *Q*_*ϕ*_(**z**|**x**) is a multivariate Gaussian distribution

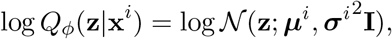

where ***μ***^*i*^ and ***σ***^*i*^ are outputs of encoder for **x**^*i*^. The VAE further uses the reparameterization trick **z**^*i,l*^ = ***μ***^*i*^ + ***σ***^*i*^ *⊙* ***ϵ***^*l*^, ***ϵ***^*l*^ *∼ 𝒩* (**0, I**), where *⊙* means element-wise product, to sample from the posterior *Q*_*ϕ*_(**z**|**x**^*i*^). By sampling **z**^*i*^ *L* times, the ELBO of the model can then be estimated as:

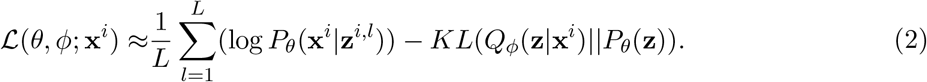

The VAE makes use of two neural networks as encoder and decoder, and applies stochastic gradient descent to optimize the objective. The encoder network with parameters *ϕ* takes **x**^*i*^ as input and outputs the parameters (***μ*** and ***σ***) in the approximate posterior *Q*_*ϕ*_(**z** | **x**). By generating **z**^*i*^ from this posterior distribution as an input, the decoder network with parameters *θ* outputs the parameters in the conditional distribution *P*_*θ*_(**x**|**z**).

### 3.2 CoupleVAE for perturbational single cell RNA-sequencing data prediction

CoupleVAE is developed under the framework of VAE. It aims to generate gene expression data for perturbed cells from that for controlled cells. It consists of three modules: inference process(encoder), coupling process(coupler), and generative process(decoder). The details for the inference process, generative process, and variational lower bound are described as follows.

#### 3.2.1 Inference Process

CoupleVAE contains two recognition models in the inference process, which independently produce the latent variables **z**_*c*_ and **z**_*p*_ from the input **x**_*c*_ and **x**_*p*_, respectively, The true posteriors of **z**_*c*_ and **z**_*p*_ are denoted as *P*_*θ*_(**z**_*c*_|**x**_*c*_) and *P*_*θ*_(**z**_*p*_|**x**_*p*_), respectively. Since it is intractable to calculate the integral of posterior as in VAE, we introduce a distribution *Q* parameterized by *ϕ* to approximate the true posterior. Due to the independence of the two recognition models, the approximate posterior can be written as

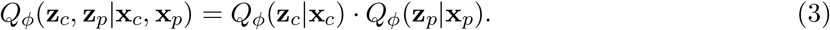

In practice, the CoupleVAE takes the prior over the latent variables **z**_*c*_ and **z**_*p*_ as *P*_*θ*_(**z**_*c*_) = 𝒩 (**0, I**) and *P*_*θ*_(**z**_*p*_) = 𝒩 (**0, I**), and assumes approximate posterior distribution to be multivariate Gaussians

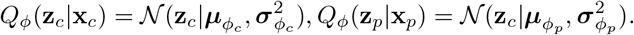

#### 3.2.2 Coupling Process

CoupleVAE assumes a coupling process of two hidden variables in the latent space, that is, latent variable **z**_*c*_ and **z**_*p*_ are transformed to **z**_*p*_ and **z**_*c*_ via conditional distributions 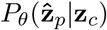 and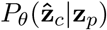, respectively. It realizes the mutual translation between the controlled cell and the perturbed cell in the latent space by forcing 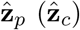 and **z**_*p*_ (**z**_*c*_) to be close. This connects the inference process and generative process.

By assuming the independence between the two coupling distributions, as shown in Figure 1, we could have

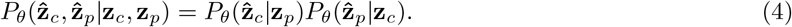

In practice, CoupleVAE takes the coupling distributions as multivariate Gaussians:

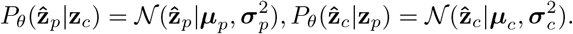

#### 3.2.3 Generative Process

CoupleVAE includes two generative processes. One is to reconstruct **x**_*c*_ and **x**_*p*_ from **z**_*c*_ and **z**_*p*_ respectively, which is the same as VAE, and the other is to reconstruct **x**_*c*_ and **x**_*p*_ from hidden variables 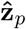 and 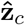, respectively. As shown in Figure 1, CoupleVAE generates **x**_*p*_ for the perturbed cell from the translated latent vector 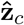 through 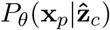, while generates **x**_*c*_ for the controlled cell from the translated latent vector 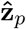 through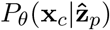.

By assuming the independence of the generative process for **x**_*p*_ and **x**_*c*_, which is also shown in the graph model in Figure 1, we could have generative model *P*_*θ*_(**x**_*c*_| **z**_*p*_), *P*_*θ*_(**x**_*p*_ |**z**_*c*_) (reconstruction) and 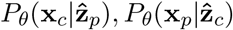 (transform-reconstruction). In practice, the CoupleVAE takes the generative probability as

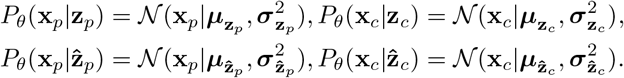

#### 3.2.4 Variational Lower Bound

To learn the inference, coupling and generative processes based on training pairs, the objective of CoupleVAE is to maximize a sum over the log likelihood of the pairs of cells (**x**_*c*_, **x**_*p*_) in the training set, i.e. Σ log *P*_*θ*_(**x**_*c*_, **x**_*p*_). We theoretically prove that the log likelihood of joint distribution for (**x**_*c*_, **x**_*p*_) has the following lower bound:

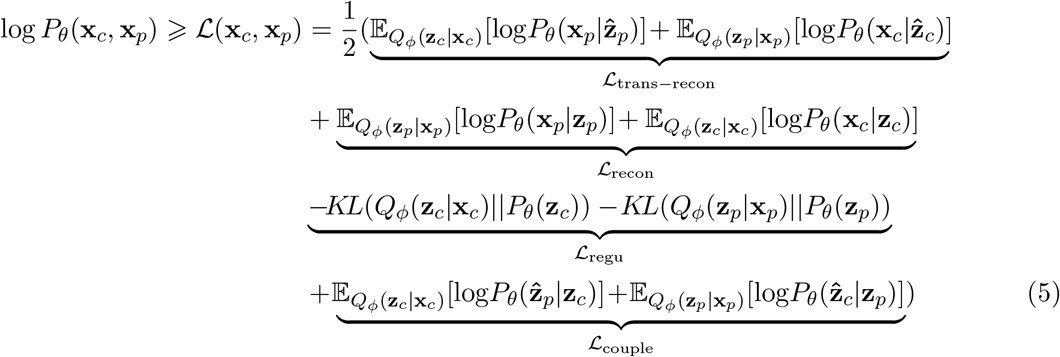

From the equation (5), the ELBO consisting of four items is formulated as:

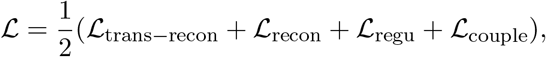

where reconstruction errors *ℒ*_trans*−*recon_ and *ℒ*_recon_ are to minimize the similarities between the generated cells and the true cells, regularization term *ℒ*_regu_ is to ensure the closeness between posterior distributions of **z**_*p*_ and **z**_*c*_ and the prior distributions, and the coupling loss *ℒ*_couple_ enables the mutual translation in the latent space, i.e., the transformed **z**_*c*_ (or transformed **z**_*p*_) is similar with **z**_*p*_ (or **z**_*c*_) in the latent space. More details of the theoretical and empirical loss function are shown in Supplementary Note 5.

### 3.3 Model training

Our experiments are divided into two setups, independent-and-identically distributed (i.i.d.) [26] and out-of-distribution (o.o.d.) [22]. The COVID-19 and PBMC datasets use the i.i.d. setup and the LPS6 dataset uses the o.o.d. setup. In the i.i.d. setup, the training set, validation set, and test set are then divided according to a ratio of 8:1:1. In the training process, CoupleVAE is fed cells of the controlled and perturbed states from one specific cell type. In the latent space, training of the model constructs the transformation from the cells in the control state to the cells in the perturbed state, resulting in simulated cells being more similar to the cells in the perturbed state. For prediction, we input the controlled cells from the test set into the encoder, which is then transformed into a latent vector that is perturbed. After going through the decoder, the output is the predicted data. In the o.o.d. setup, the model cannot access cells in holdout condition. When we predict a perturbation for a particular cell type, the training data are all cell types except the predicted one. In the cross-species prediction task on the LPS6 dataset, we have two training strategies corresponding to the two training models as condition-CoupleVAE and species-CoupleVAE. For condition-CoupleVAE, we use the cells from all species in the control state of the training set (red box in Fig. 4a) as **x**_*c*_, and the cells from all the species in the LPS state as **x**_*p*_. For species-CoupleVAE, we use mouse cells in the training set (red box in Fig. 4e) as **x**_*c*_ and rabbit cells as **x**_*p*_. We input the mouse cells in the LPS state as the test data into species-CoupleVAE, and finally get the rabbit cells in the LPS state. In all settings, we keep the same number of cells for each cell type in two conditions.

### 3.4 Single-cell RNA-seq datasets

- **The COVID-19 dataset**. The COVID-19 dataset and its metadata are available for download from [29]. We collected the processed dataset from [25]^1^. The dataset was filtered for cells with at least 500 expressed genes and genes expressed in at least 5 cells. Cell counts were normalized, and the top 6000 highly variable genes were selected. Finally, we log-transformed the data. 11742 cells and six cell types are included after data pre-processing, which contains 402 CD8-T cells, 5334 macrophages, 1340 NK cells, 3206 neutrophil, 792 secretory and 668 mDC. They are derived from a combination of data from lung [38] and bone marrow [39], [40] and COVID-19 samples.
- **The PBMC dataset**. This dataset is a single-cell gene expression count dataset of peripheral blood mononuclear cells (PBMCs), which is used to generate a reference transcriptome profile [41]. The investigators annotate the cell types by extracting the average top 20 clustered genes for 7 of the 34 identified cell types in the PMBCs dataset. We collected data from the work [21] ^2^ and processed the expression data to retain the top 6,998 highly variable genes and 12,982 cells. This dataset contains 1414 B cells, 4874 CD4-T cells, 974 CD8-T cells, 1062 CD14-Mono cells, 814 DC cells, 1934 FCGR3A-Mono cells and 924 NK cells.
- **The LPS6 dataset**. The LPS data set (accession id E-MTAB-6754) was obtained from BioStudies [35]^3^. The data were further filtered for cells, normalized and log-transformed. We used BiomaRt (v.84) to find ENSEMBL IDs of one-to-one orthologs in the other three species with mouse. A total of 6,619 genes were selected from all species for training the model. The final data includes 77,642 cells. For more preprocessing information about this dataset, please refer to the paper on scGen [21]. The characteristic of this dataset is that it contains different species, and the distribution between two groups of each species is quite different. It is mainly used to explore the performance of the models when predicting single-cell perturbation across species.

### 3.5 Comparison methods

We choose as benchmarks the previously well-performing single-cell perturbation prediction models, which are scPreGAN, trVAE, scGen, and CVAE.

- **scPreGAN** [20]. It is a deep generative model for predicting the response of single-cell expression to perturbation. This model combines the architecture of AE and GAN, the former is to extract common information of the unperturbed data and the perturbed data, the latter is to predict the perturbed data.
- **trVAE** [22]. It is an MMD-regularized conditional VAE, that matches distributions across conditions using maximum mean discrepancy in the decoder layer that follows the bottleneck.
- **scGen** [21]. It is a model combining VAE and latent space vector arithmetics for high-dimensional single-cell gene expression data. scGen simulates cell type, study and species responses to perturbation and infection, enabling learning of cell type and species-specific responses.
- **Standard CVAE** [23]. The Conditional Variational Autoencoder (CVAE) is an extension of the Variational Autoencoder (VAE) that introduces conditional variables to improve the generation process. In the CVAE setting one can train a model conditioned on two existing biological conditions.

For each method, we use the default parameters for training.

### 3.6 Evaluation metric

The Pearson correlation coefficient between two variables is defined as the quotient of the covariance and standard deviation between the two variables. It is obtained by estimating the covariance and standard deviation of the sample:

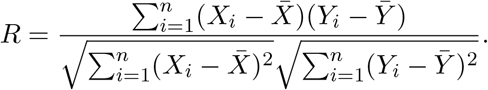

The squared Pearson correlation coefficient *R*^2^ varies from 0 to 1. All the *R*^2^ values are calculated by squaring the *rvalue* output of the *scipy*.*stats*.*linregress* function and denote squared Pearson correlation.

## 4. Conclusion

In this work, we have introduced CoupleVAE, a novel deep generative model, for predicting single cell responses to perturbation measured by scRNA-seq data. CoupleVAE consists of two encoders, a coupler and two decoders, which do not share parameters between them. We devised the transformations between the latent variables in the latent space such that the predicted cells can be as close to the real perturbational cells as possible. The existing VAE-based models do not perform well in perturbation response prediction. It is most likely because of ignorance of the heterogeneity in different cell types’ responses to perturbations. Through applying two separate encoders to obtain the latent vectors for different conditions, and constructing the transformations between them, the effects caused by different perturbations are learned well in a low-dimensional representation of gene expression. Extensive experiments on PBMC and COVID-19 datasets demonstrate the effectiveness of CoupleVAE in the perspective of mean gene expression prediction, common DEGs detection and common response markers expression prediction.

Although CoupleVAE works well for most cases, it is far from perfect, for example, when predicting NK cells in the PBMC dataset and the COVID-19 dataset. This may be due to the small sample size of NK cells, or the complex role of NK cells in the immune system. For example, it has been shown that the severity of COVID-19 disease is associated with reduced NK cell numbers, NK cell exhaustion and lack of certain mature effective NK cell phenotypes [42, 43]. The mapping of interconversions between conditions in latent space will be designed to be more complex in order to capture complex perturbation response information. In addition, cross-species perturbation prediction is an interesting direction of research, since there are still many species with differential response mechanisms among them that are not yet known. This brings a great challenge to the prediction problem.

## 5. Acknowledgements

This work was supported by National Natural Science Foundation of China project under Grant No. 12222115.

## Code availability

The software is available at https://github.com/LiminLi-xjtu/CoupleVAE.

## Supplementary Materials

### Supplemental Note 1

In the cross-species experiment, we investigate the differences between different states of the same species(Fig. 4a)and between species in the same state(Fig. 4e, Supplementary Fig.12), and between different species in different states(Fig. 4). The ultimate goal of this experiment is to predict the gene expression values of unknown perturbed species cells, e.g., we remove the mouse cells in the dataset and use the remaining cells as a training set to predict the perturbed mouse cells. A deep understanding of the differences between species and between states is essential for experiments. CoupleVAE trained two models, condition-CoupleVAE and species-CoupleVAE(see Fig. 4), in predicting the gene expression values of each perturbed species. That is, in training condition-CoupleVAE, we use unperturbed rabbit, rat and pig cells as **x**_*c*_ in the model, and perturbed rabbit, rat and pig cells as **x**_*p*_ in the model. When training the species-CoupleVAE, we use unperturbed rabbit cells as **x**_*c*_ and unperturbed mouse cells as **x**_*p*_. In the condition-CoupleVAE prediction stage, we input unperturbed mouse cells into condition-CoupleVAE to get the predicted perturbed mouse cells. In the species-CoupleVAE prediction stage, we input the perturbed rabbit cells into the condition-CoupleVAE to get the predicted perturbed mouse cells. Our model deeply portrays the differences between differences between species and between states. For the other models, they only roughly describe the differences in the states between species, and the differences between species and states are not explicitly described. For example, scGen simply inscribes the differences between species and states as two vectors in different directions in the hidden space, and then adds and subtracts the feature representations to get the final result. Whereas trVAE just closes the distance between the distributions of different states for subsequent transformations and does not explain deeper differences.

### Supplemental Note 2

Here, in this section, we present all of the hyperparameters of the proposed scGen, trVAE, scPreGAN and CVAE used in the paper. We used early-stopping criterion for scGen, trVAE and CVAE: the training stopped after 20 consecutive epochs with improvement less than a threshold on the validation loss. All architectures use dropout regularization, batch normalization and Adam optimizer. We use a *L*_2_ regularization for covid-19, PBMC, and LPS6 datasets using a scale of 0.1.

### Supplemental Note 3

In plotting a bar graph representing the Pearson correlation coefficient, our experimental setup is to subsample each cell type 100 times, with 80% of the total number of cells randomly selected for each sampling. The Pearson correlation coefficients are calculated for the cells they sampled, and the numbers on each bar in the plot represent the average of the one hundred samples, error bars depict s.d. For the selection of differentially expressed genes, we use *scanpy*.*tl*.*rank genes groups* from the scanpy library, and the method uses Wilcoxon rank-sum (Wilcoxon).

### Supplemental Note 4

Often, the conditions are not equal in size, leading to biased reconstructions. Imbalanced data refers to a situation where the distribution of classes in a dataset is not equal. In other words, one class has significantly more instances than the other class. This is a common problem in machine learning and can occur in various domains such as fraud detection, medical diagnosis, and text classification. Imbalanced data can cause several problems in machine learning, such as inaccurate error evaluation. To address the imbalanced learning problem, we undersample the majority class examples. [44] To prevent similar problems, we balanced cell type(species) and condition size before training.

### Supplemental Note 5

The loss function of CoupleVAE has a term coupling loss, which is designed to allow the two conditional data distributions to be transformed into each other in the latent space. It has been proved that this term plays an important role. For the LPS dataset, we use the latent space representation of condition-CoupleVAE. As shown in the Supplementary Figure 13, the transformed distributions of all four species overlap exactly with the real data distributions of the four species that are perturbed by the LPS. In contrast, the latent space expression of the data predicted by scGen overlaps with the gene expression in the control state.

### Supplemental Note 6

We can show that the log likelihood of each joint distribution of (**x**_*c*_, **x**_*p*_) has the following lower bound:

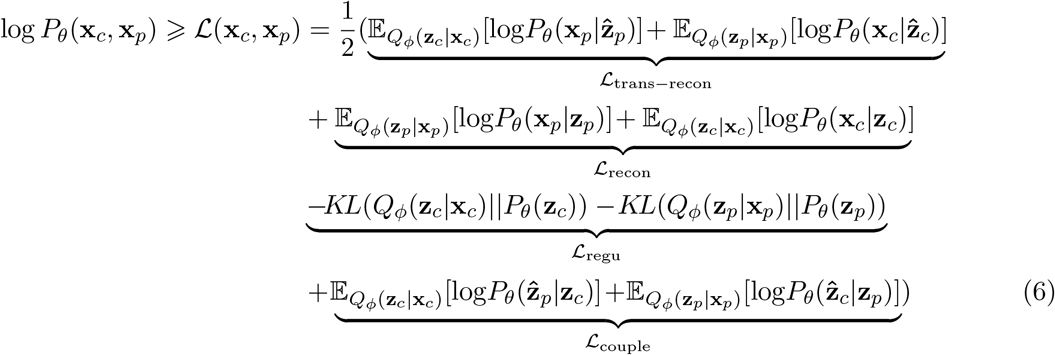

To show the upper bound, we could first obtain the ELBO by

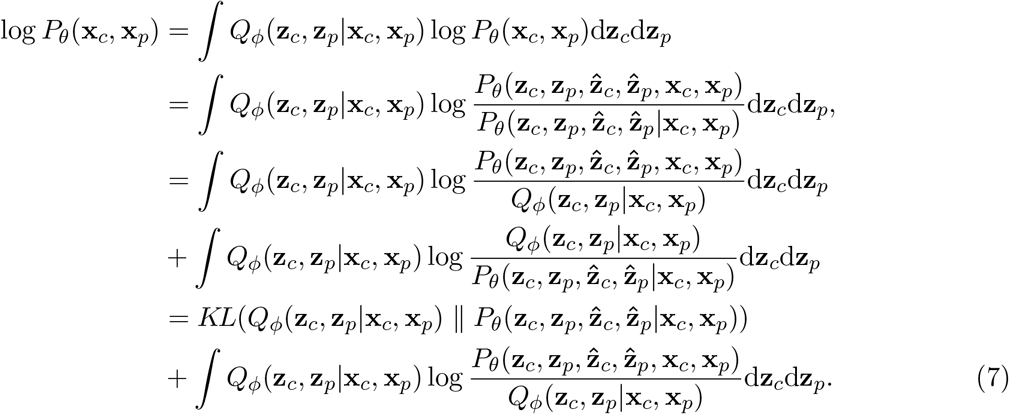

To further simplify the equation (7) by the independence assumption in equation (4), which is also shown in the graph model in Figure 1, the joint distribution 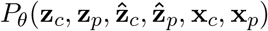 can be expressed as

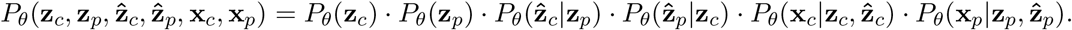

To obtain the decompositions of 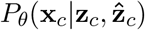 and 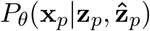, we perform a simple derivation. Suppose there are random variables *a, b* and *c*, where *b* and *c* are independent. We have

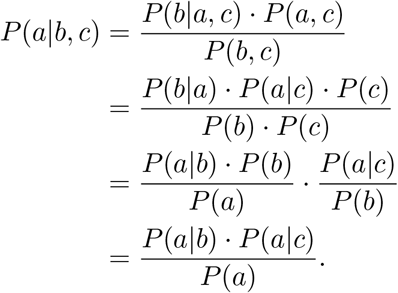

The expression for the joint probability distribution is eventually written as

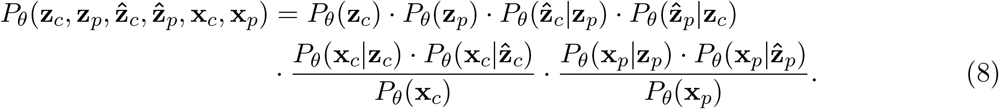

In equation (7), we denote the KL term as *D*_*KL*_. And the numerator in equation (8) is denoted as *M*.

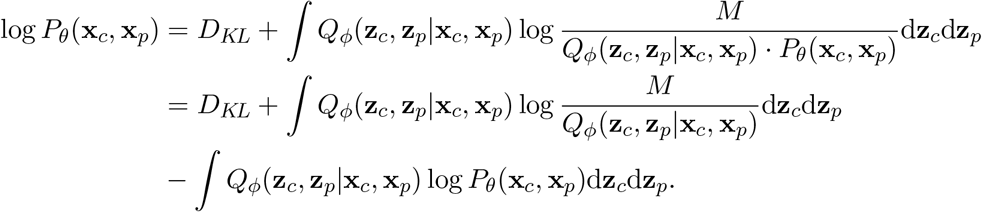

Finally, we have

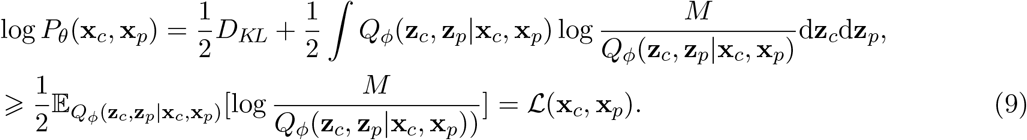

By equations (3) and (9), the ELBO is further simplified as follow:

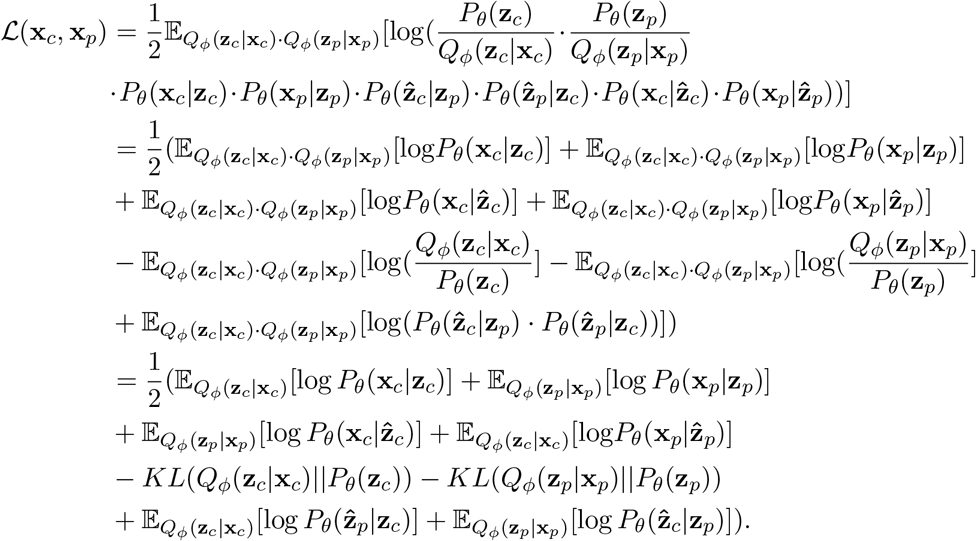

We jointly optimize the generative model *θ* and variational *ϕ* parameters to maximize the ELBO of the CoupleVAE for the training data.

ℒ_recon_ and ℒ _couple_ are estimated by Monte Carlo sampling. Using the reparametrization trick, we can obtain an unbiased gradient estimator of the distribution parameters by backpropagation, which is known as Stochastic Gradient Variational Bayes (SGVB [36]). Thus we have

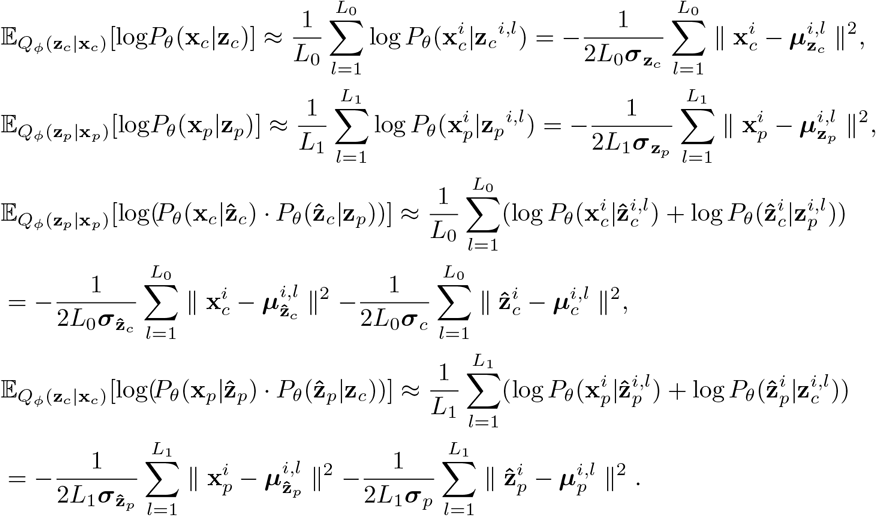

And the ℒ _regu_ do not need to be approximated because the kl divergence between two normal distributions has an explicit solution. Finally, we calculate the approximate gradients of *θ* and *ϕ* and update the parameters with Adam [45]. The objective function to maximize in the CoupleVAE at 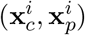 is:

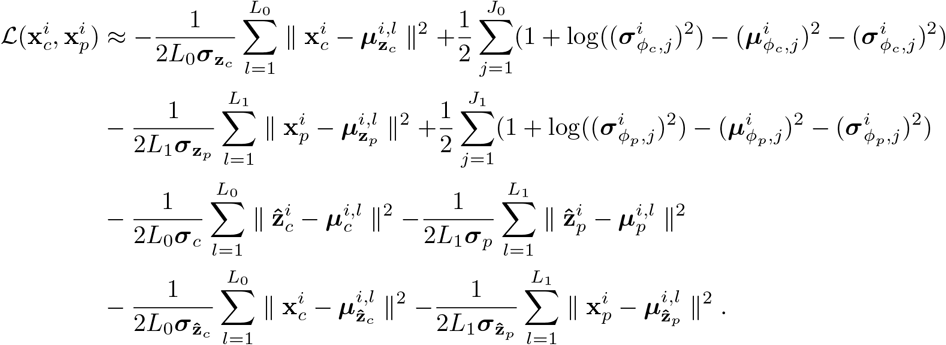

**Supplementary Figure 1:**
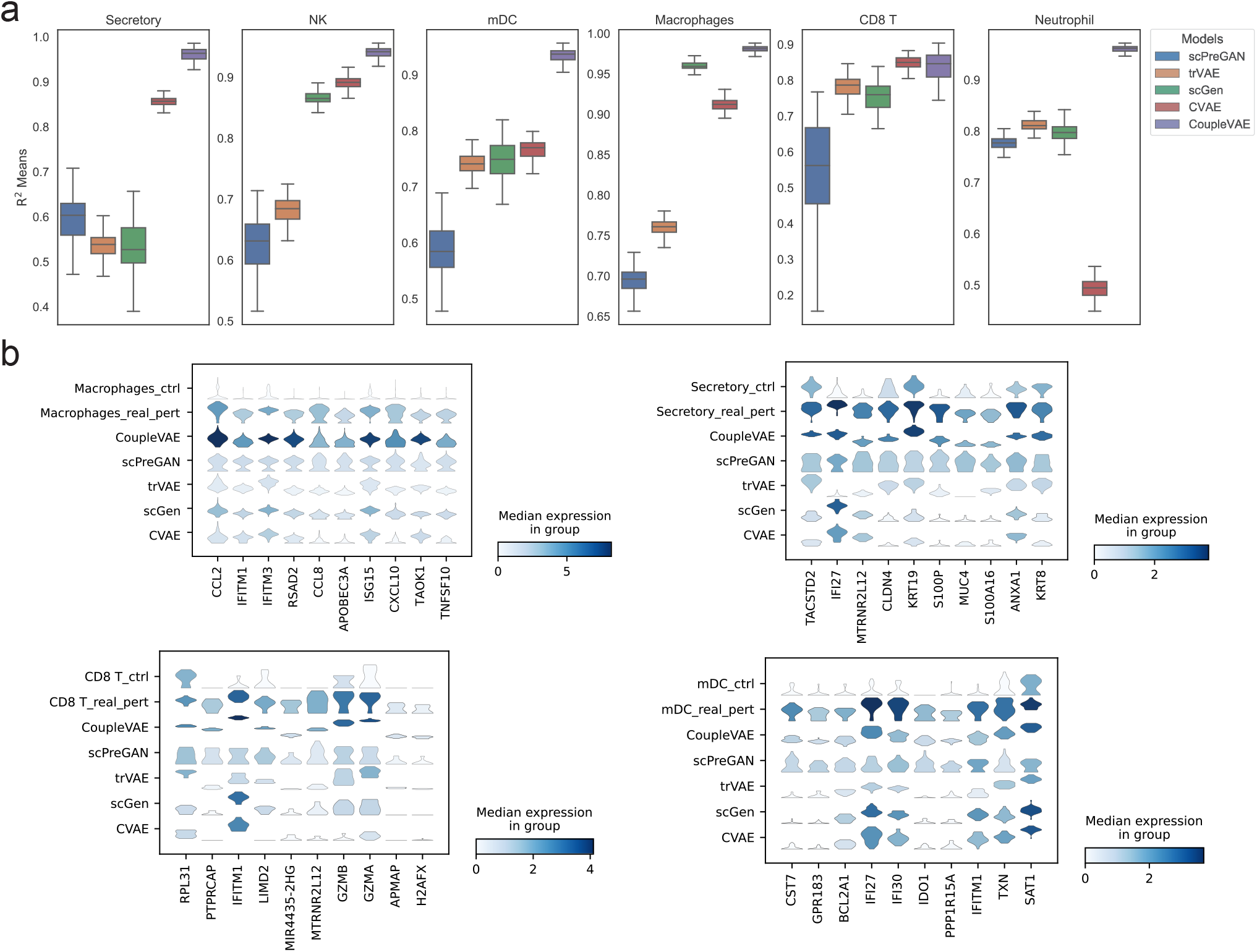
**a** Box plot of Pearson’s *R*^2^ values for the average expression of top 100 DEGs are compared between real and predicted cells of different models in all cell types. **b** Violin plots of 10 differentially expressed genes in other five cell types after infection by SARS-CoV-2. Vertical axis: control, true distribution of five cell types and predicted distribution of five cell types by different models. Horizontal axis: expression distribution of differentially expressed genes.

**Supplementary Figure 2:**
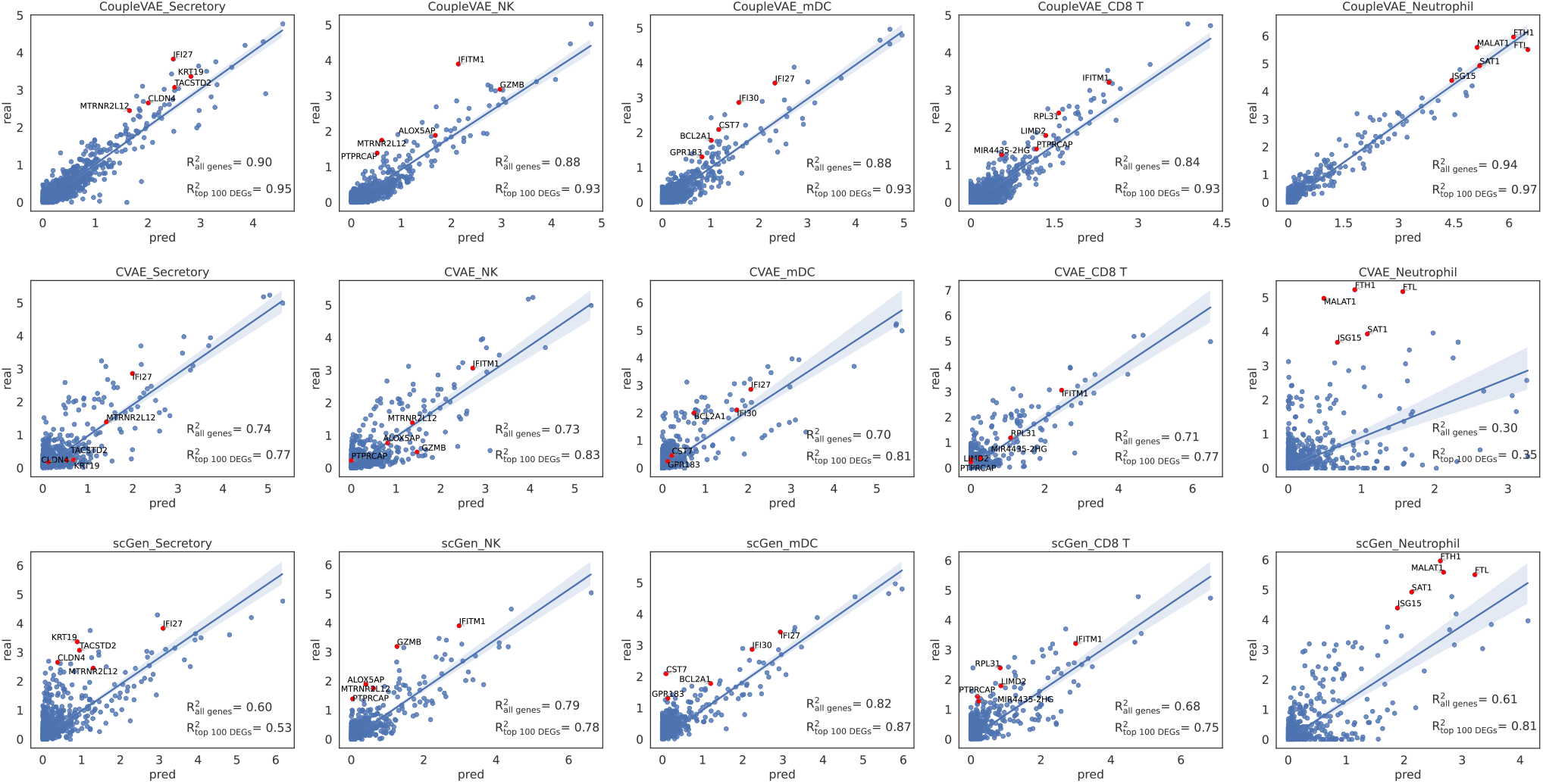
Correlation comparison between the real mean values of 6000 genes’ expression in five cell types with SARS-CoV-2 infection and those predicted by CoupleVAE, CVAE and scGen

**Supplementary Figure 3:**
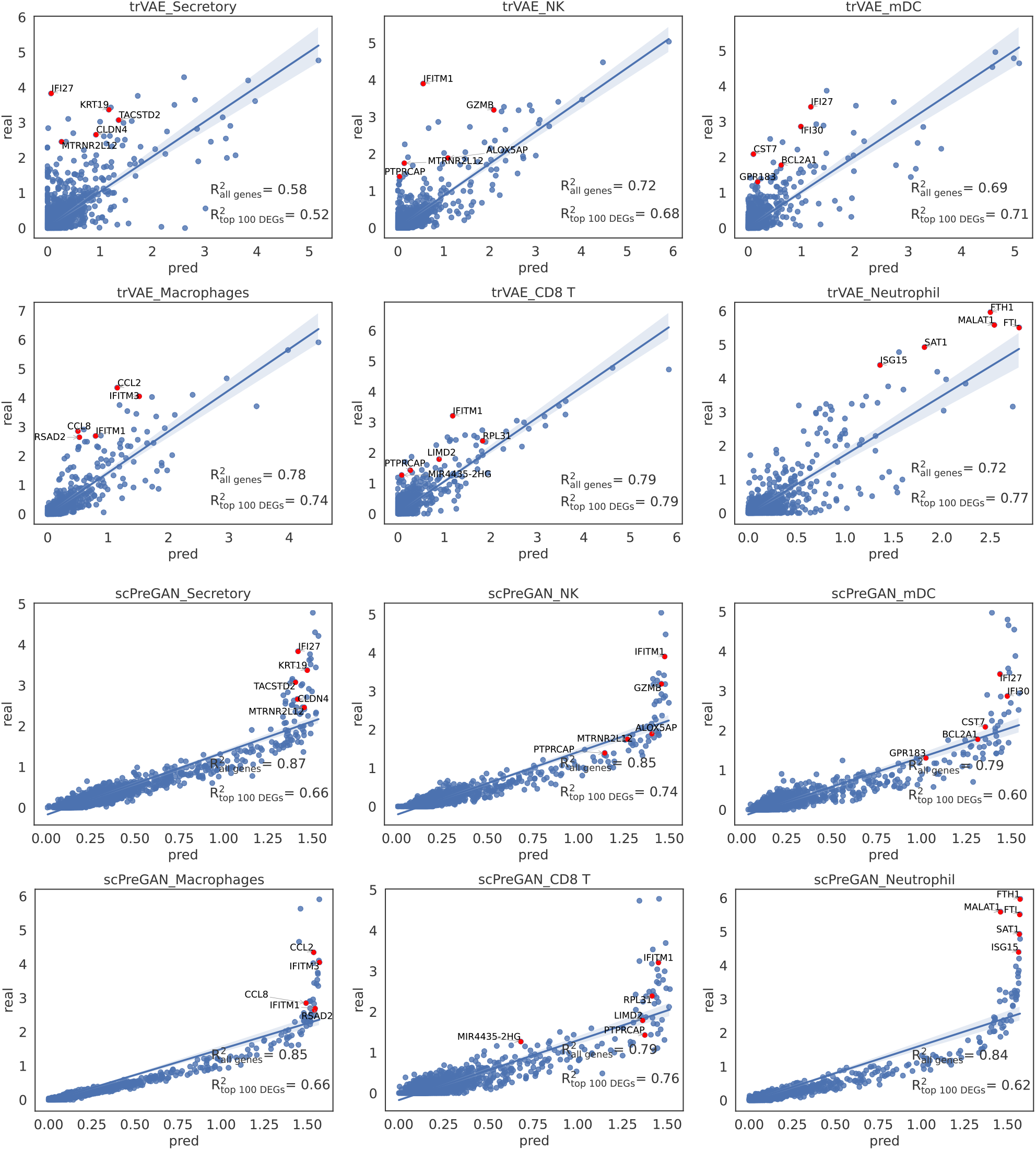
Correlation comparison between the real mean values of 6000 genes’ expression in six cell types with SARS-CoV-2 infection and those predicted by trVAE and scPreGAN

**Supplementary Figure 4:**
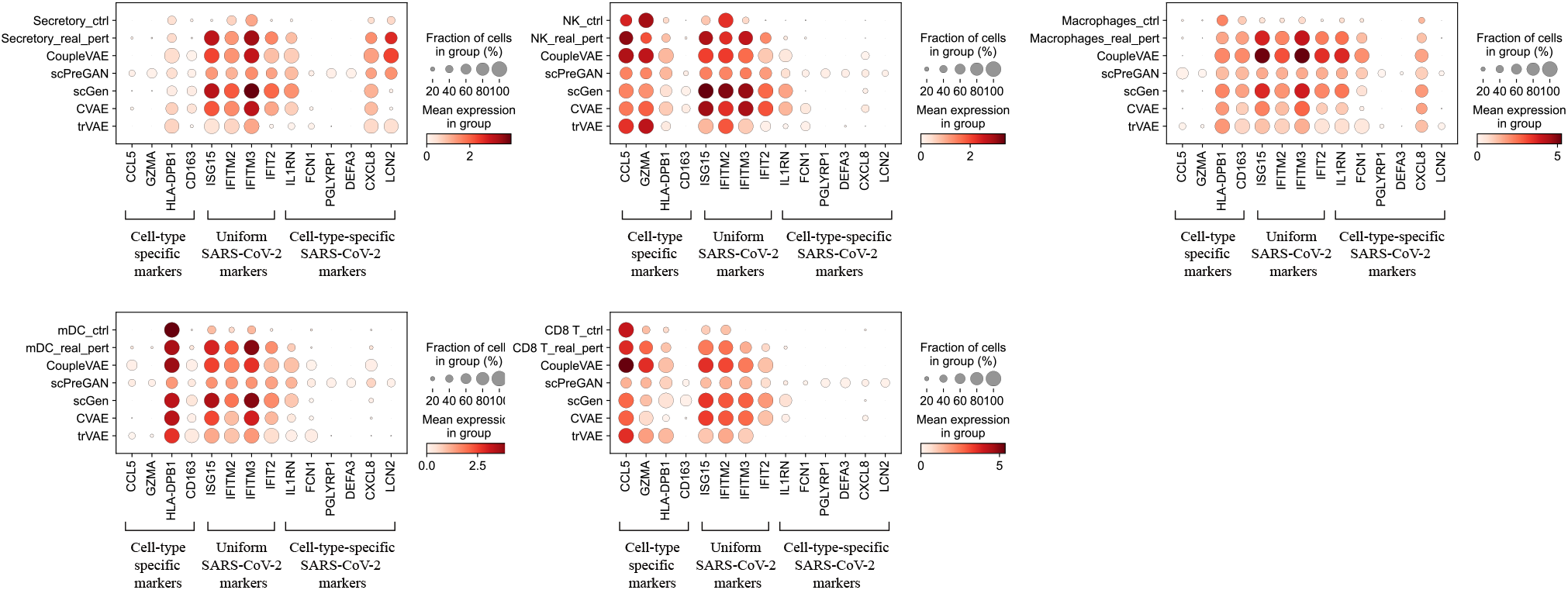
Dot plot for comparing control, predicted and real perturbation in predictions on the six cell types from COVID-19.

**Supplementary Figure 5:**
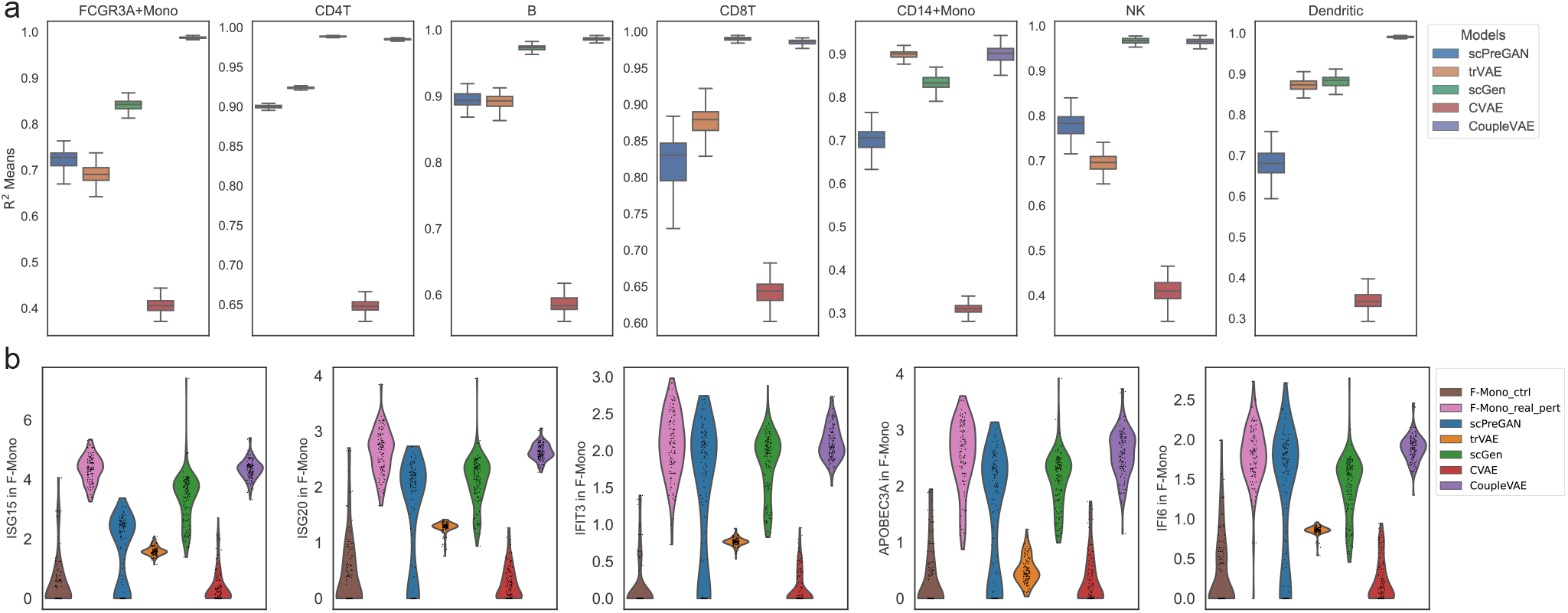
**a** Box plot of Pearson’s *R*^2^ values for the average expression of top 100 DEGs are compared between real and predicted cells of different models in all cell types. **b** Distribution of the most strongly changed genes(ISG20, IFI6, APOBEC3A, IDO1) after *IFN − β* perturbation in control, real and predicted stimulated cells of CoupleVAE compared to other prediction models in F-Mono cells.

**Supplementary Figure 6:**
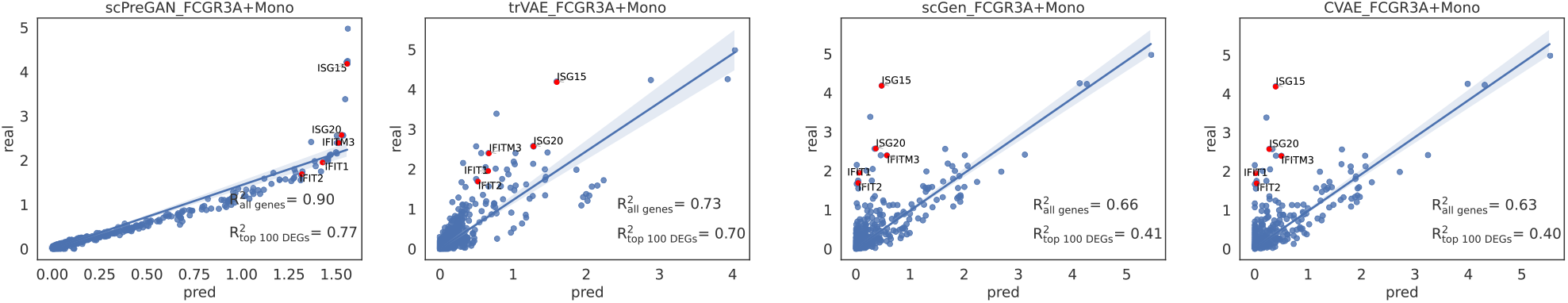
The mean value of 6998 gene expressions in 137 FCGR3A+Mono predicted using scPreGAN, trVAE, scGen and CVAE compared to the real expressions.

**Supplementary Figure 7:**
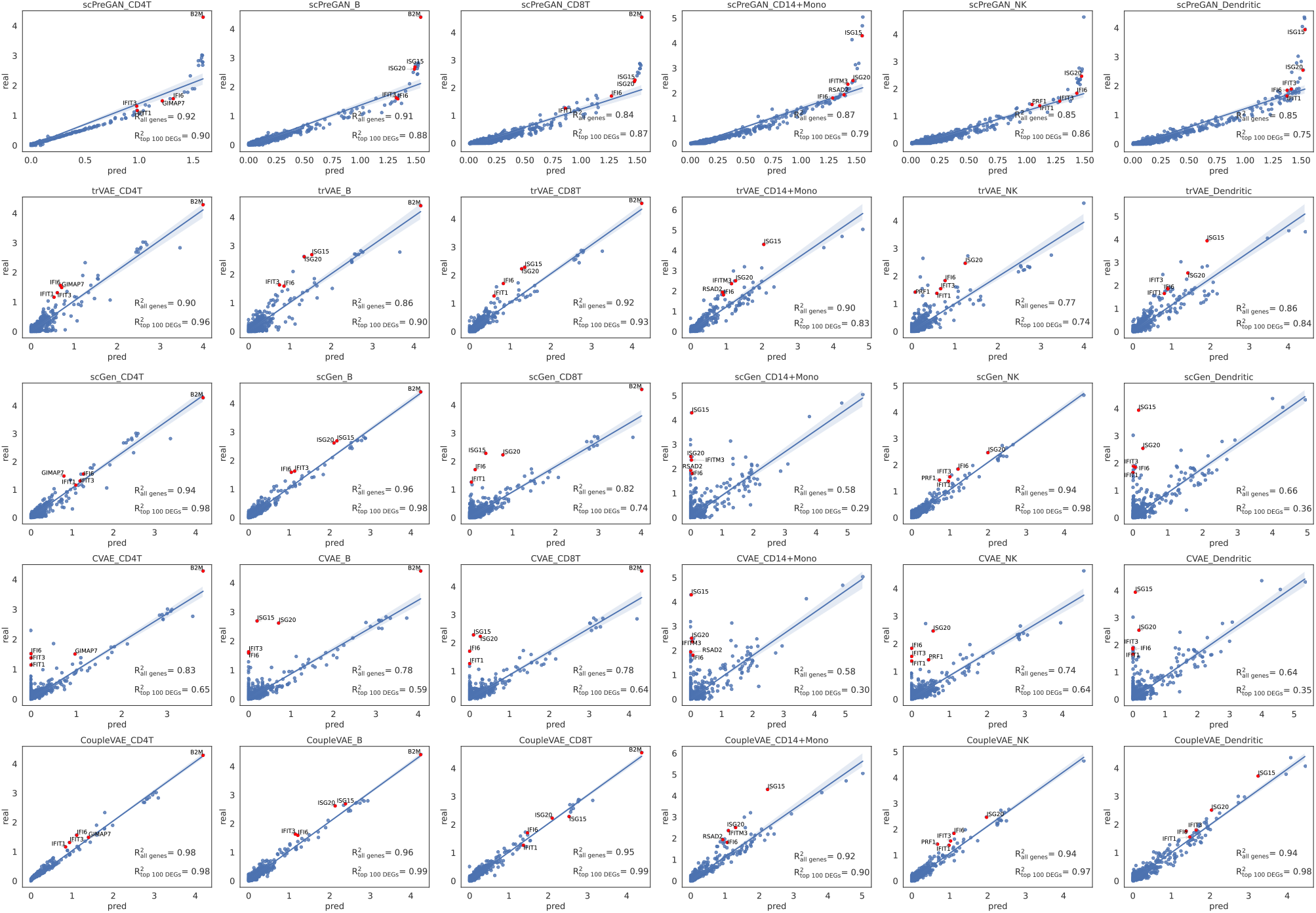
Correlation comparison between the real mean values of 6998 genes’ expression in six cell types with *IFN −β* treatment and those predicted by scPreGAN, trVAE, scGen, CVAE and CoupleVAE.

**Supplementary Figure 8:**
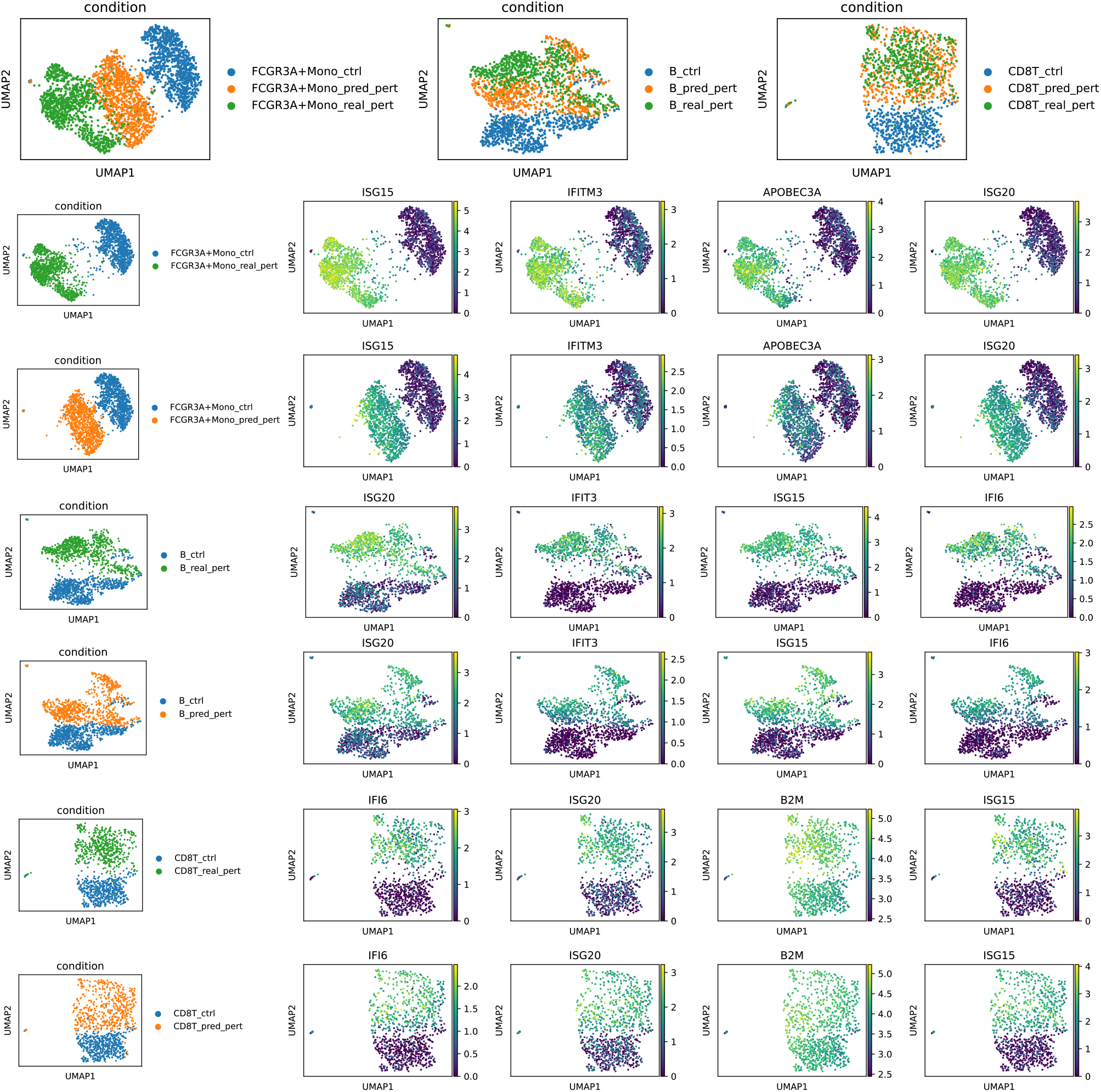
**a** The UMAP visualizations of unperturbed data, real perturbed data and predicted perturbed data for FCGR3A+Mono, B and CD8T cells. **b** The diagrams are divided into two groups, the first group being the first column and the second group being the remaining columns. The plots in the first group are UMAP visualisations of unperturbed & true perturbed data and unperturbed & predicted perturbed data for FCGR3A+Mono, B and CD8T cells. The plots in the second group are UMAP visualisations of the expression of four genes that are highly responsive to the perturbation in each cell.

**Supplementary Figure 9:**
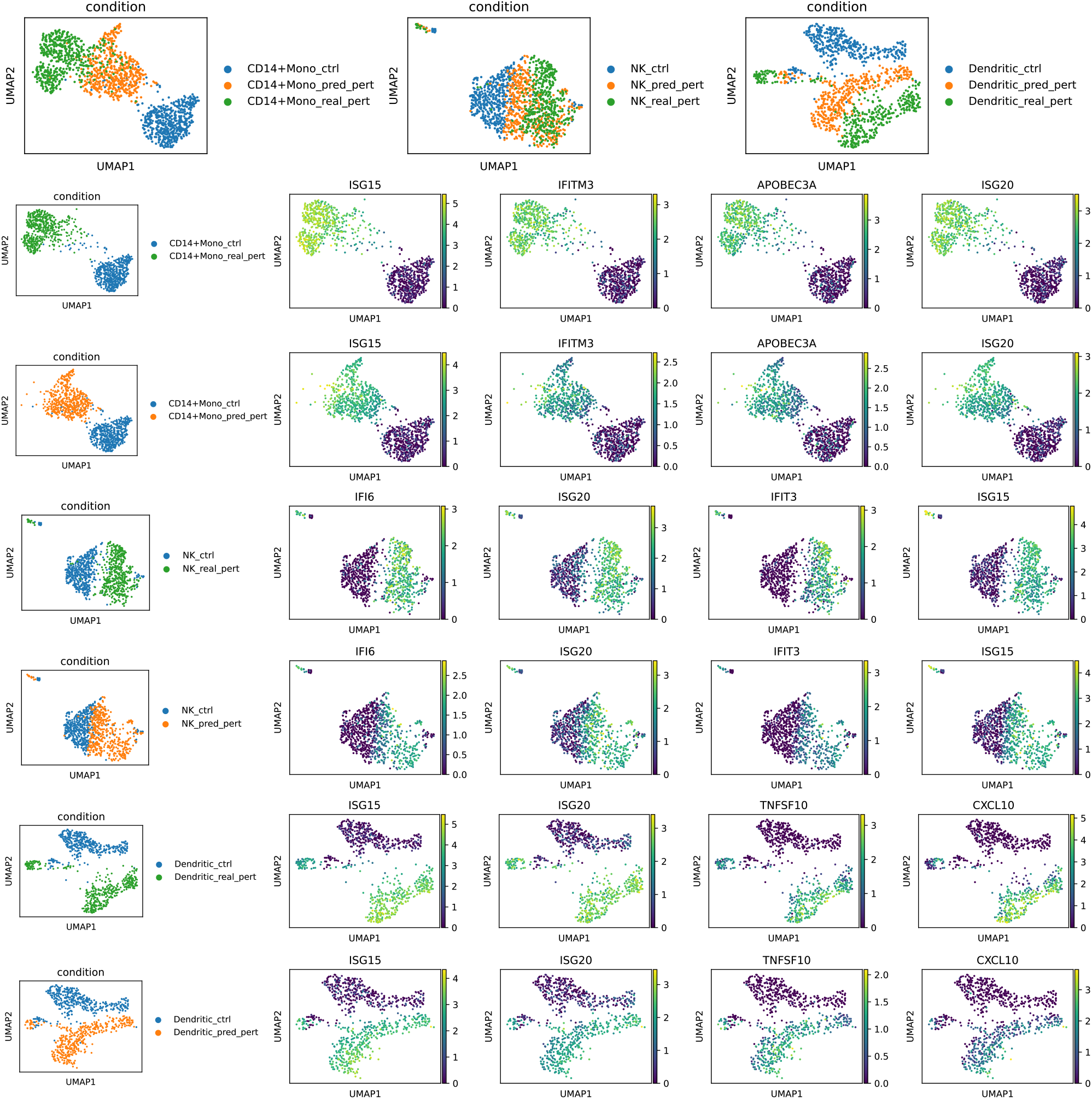
**a** The UMAP visualizations of unperturbed data, real perturbed data and predicted perturbed data for CD14+Mono, NK and Dendritic cells. **b** The diagrams are divided into two groups, the first group being the first column and the second group being the remaining columns. The plots in the first group are UMAP visualisations of unperturbed & true perturbed data and unperturbed & predicted perturbed data for CD14+Mono, NK and Dendritic cells. The plots in the second group are UMAP visualisations of the expression of four genes that are highly responsive to the perturbation in each cell.

**Supplementary Figure 10:**
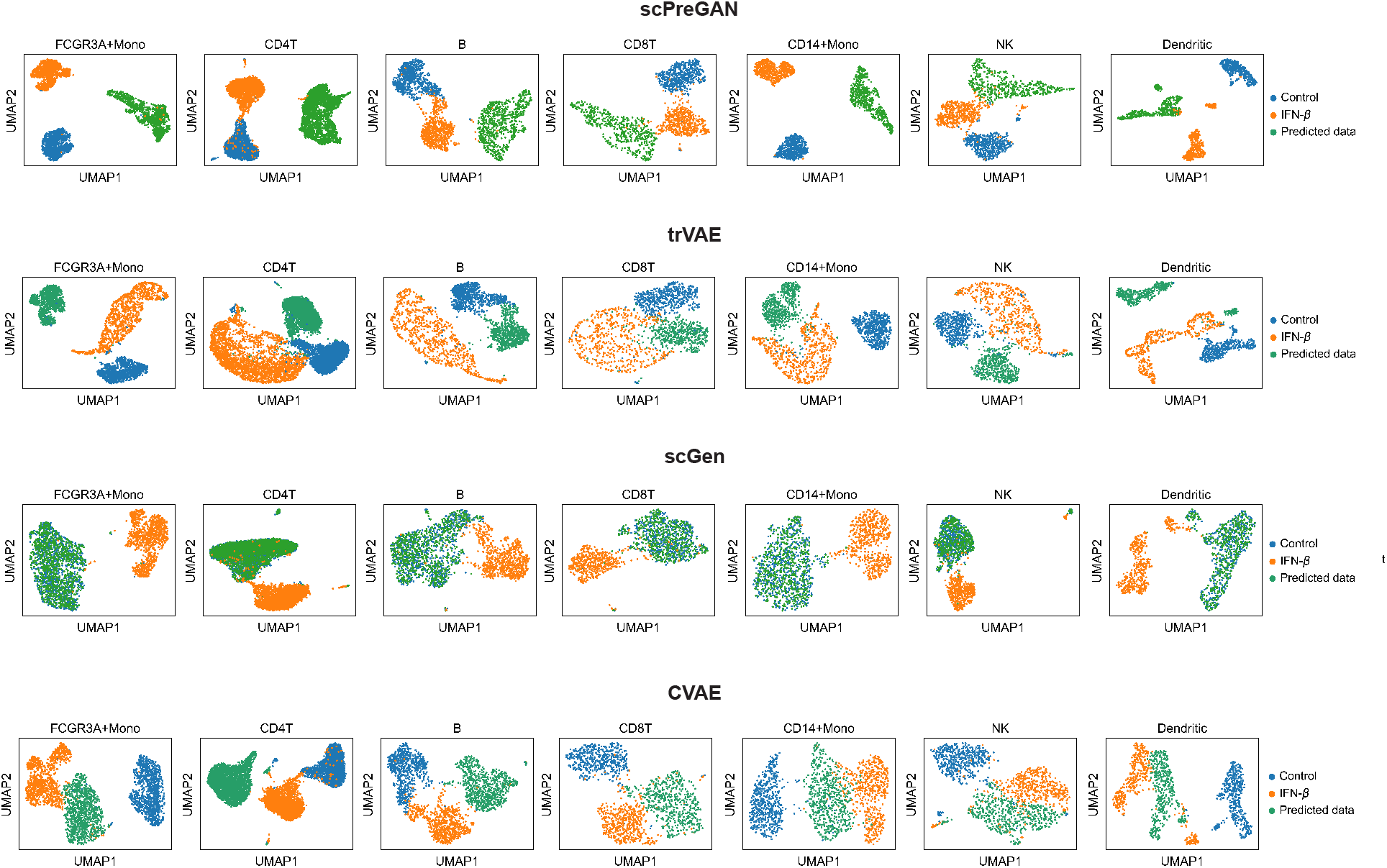
UMAP visualization of controlled data, real perturbed data and predicted data using scPreGAN, trVAE, scGen and CVAE for all seven cell types.

**Supplementary Figure 11:**
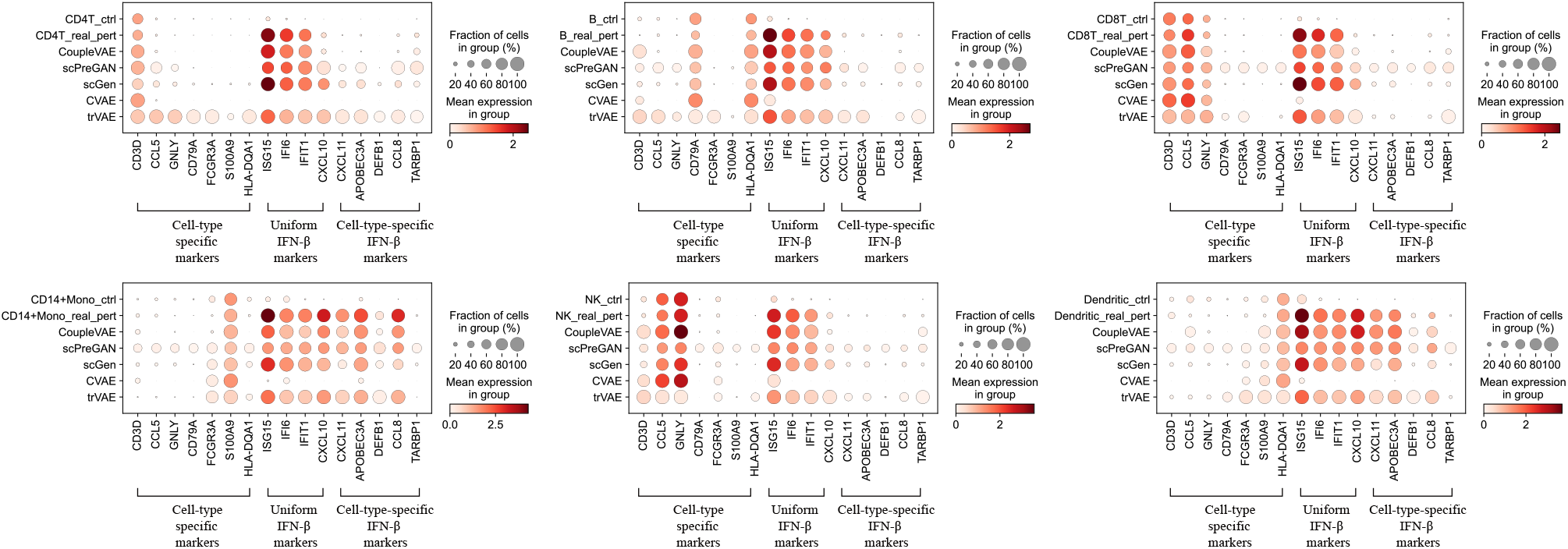
Dot plot for comparing control, predicted and real perturbation in predictions on the six cell types from PBMC.

**Supplementary Figure 12:**
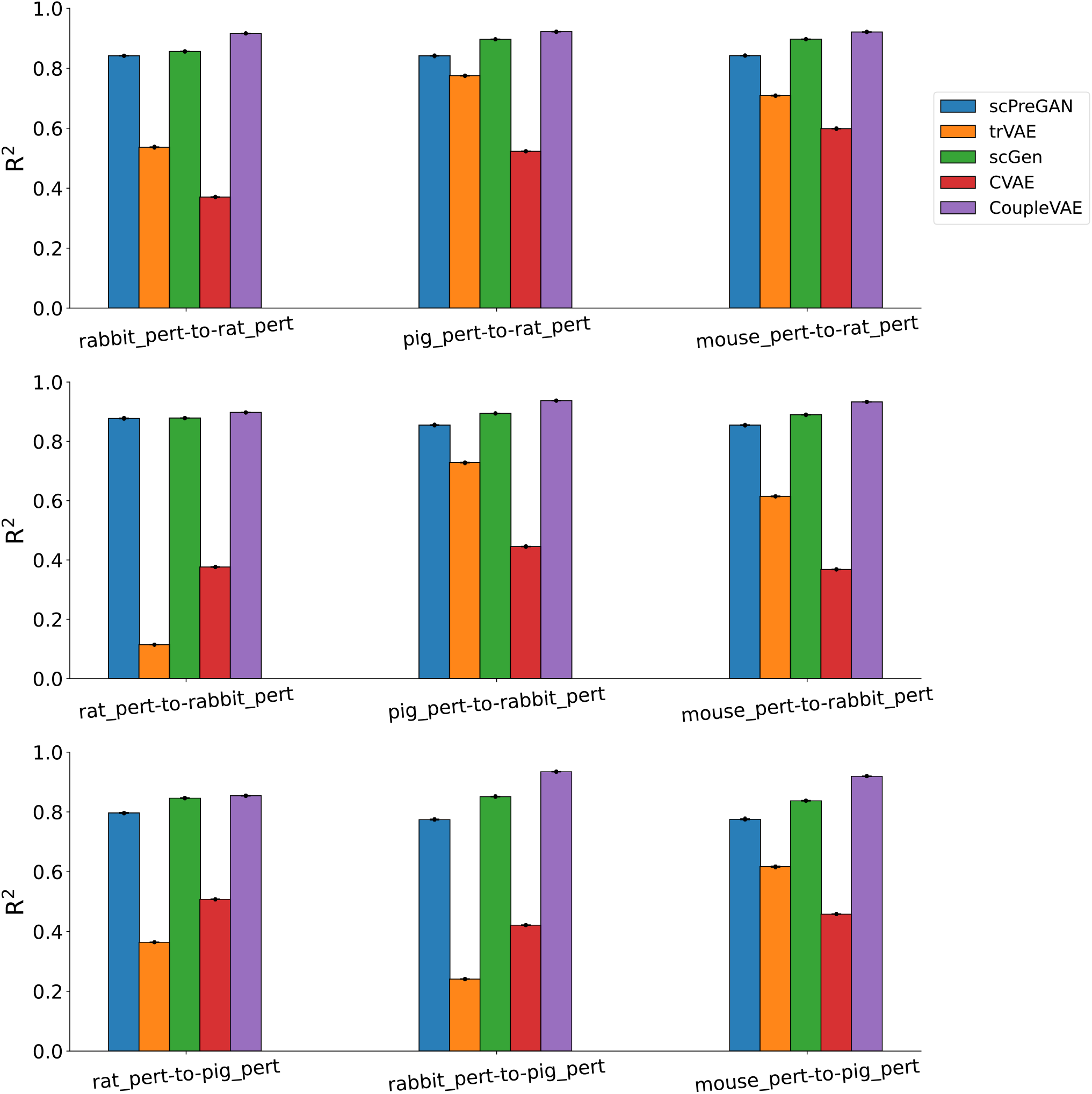
Histograms of prediction of rat, rabbit and pig cells perturbed by LPS, respectively, from the other three remaining species cells perturbed by LPS.

**Supplementary Figure 13:**
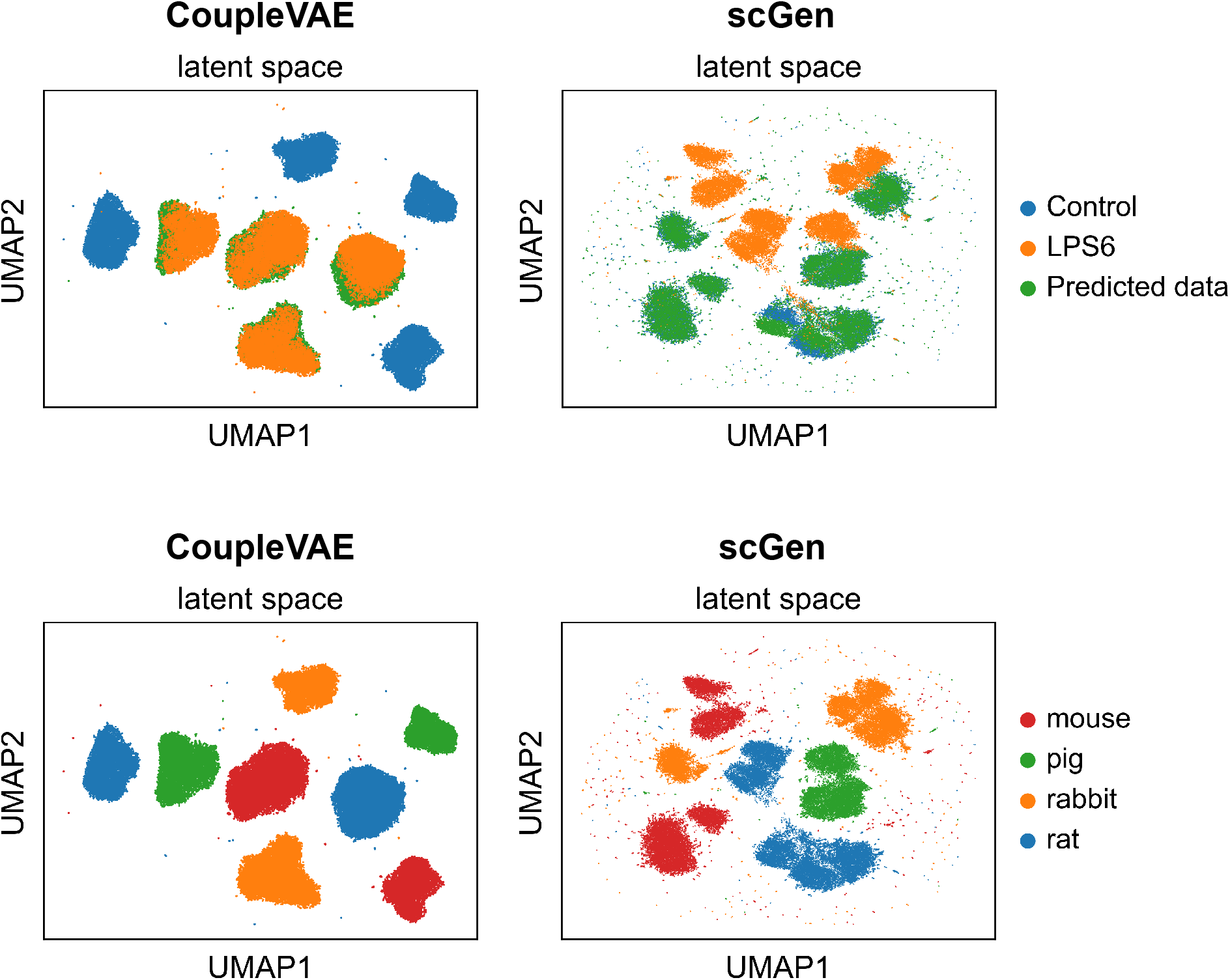
UMAP visualization of real unperturbed data, real perturbed data and predicted perturbed data for all species in latent space.

**Supplementary Figure 14:**
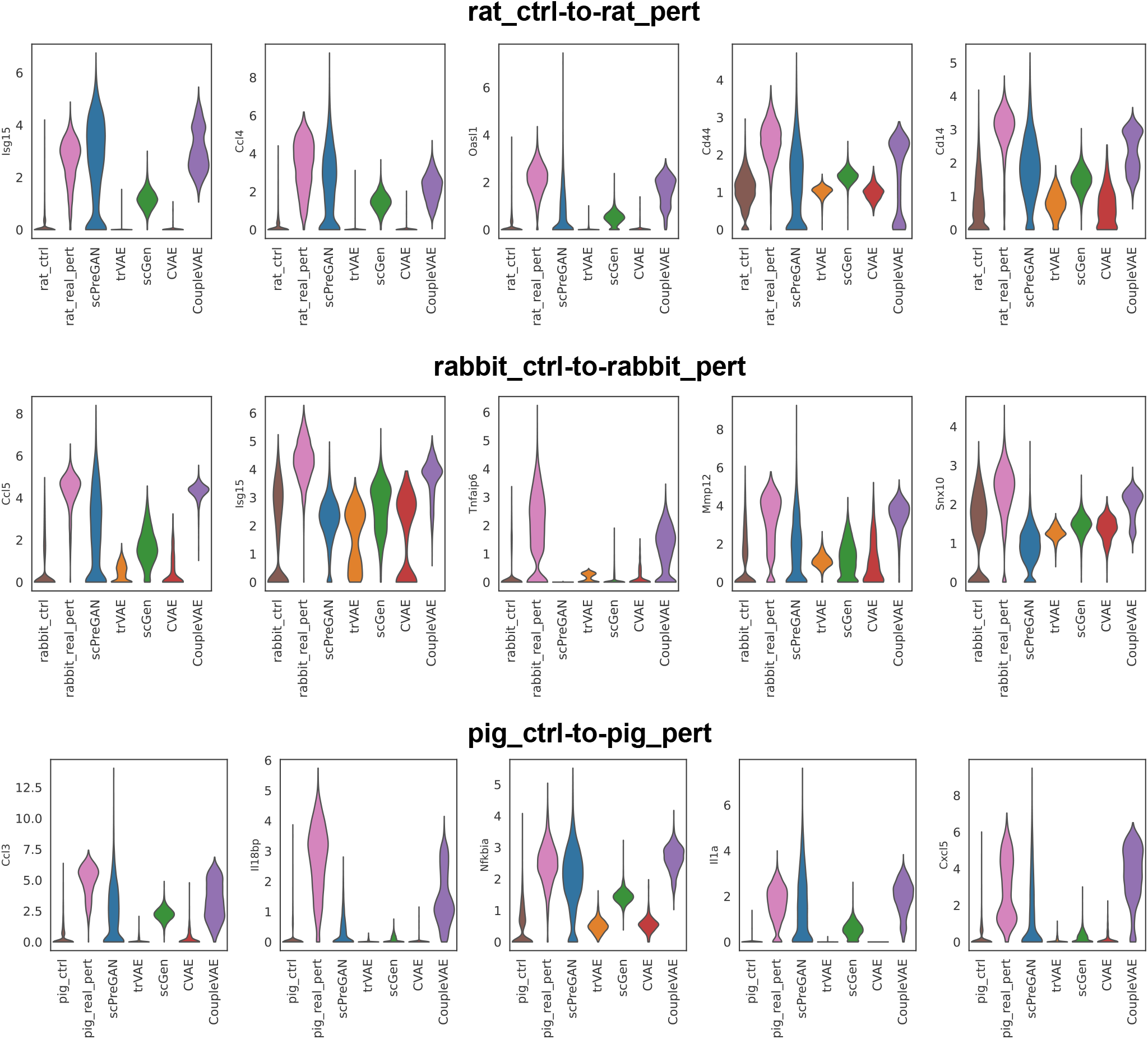
Distribution of LPS-responsive genes for rat(first row), rabbit(second row) and pig(last row) identified in the study by Haiga et al [35]. Vertical axis: expression distribution for genes. Horizontal axis: control, real and predicted distribution by different models.

## Footnotes

1 https://drive.google.com/drive/folders/1QQXDuUjKG8CTnwWWu83MDtdrBXr8Kpq

2 https://drive.google.com/drive/folders/1v3qySFECxtqWLRhRTSbfQDFqdUCAXql3

3 https://www.ebi.ac.uk/arrayexpress/experiments/E-MTAB-6754/?query=tzachi+hagai

## Notes

### Competing Interest Statement

The authors have declared no competing interest.

